# Modeling dynamic inflow effects in fMRI to quantify cerebrospinal fluid flow

**DOI:** 10.1101/2025.04.03.647027

**Authors:** Baarbod Ashenagar, Daniel E. P. Gomez, Laura D. Lewis

## Abstract

Cerebrospinal fluid (CSF) flow in the brain is tightly regulated and essential for brain health, and imaging techniques are needed to quantitatively establish the properties of this flow system. Flow-sensitive fMRI has recently emerged as a tool to measure large scale CSF flow dynamics with high sensitivity and temporal resolution; however, the measured signal is not quantitative. Here, we developed a dynamic model to simulate fMRI inflow signals based on time-varying flow velocities. We validated the model in both human and phantom data, and used it to identify important properties of the fMRI inflow signal that inform how the signal should be interpreted. Additionally, we developed a physics-based deep learning framework to invert the model, which enables direct estimation of velocity using fMRI inflow data. This work allows new quantitative information to be obtained from fMRI, which will enable neuroimaging researchers to take advantage of the high sensitivity, high temporal resolution, and wide availability of fMRI to obtain flow signals that are physically interpretable.

## 1 Introduction

The flow of cerebrospinal fluid (CSF) is essential for maintenance of brain homeostasis (Brinker et al., 2014; Johanson et al., 2008; Simon & Iliff, 2016), but the dynamics of CSF flow in the human brain remain poorly understood, in part due to technical challenges in flow imaging (Agarwal et al., 2023). CSF flow is dynamically regulated by multiple physiological mechanisms, with pulsations driven by cardiac (Feinberg & Mark, 1987), respiratory (Chen et al., 2015; Dreha-Kulaczewski et al., 2015; Kao et al., 2008) and slow hemodynamic (Fultz et al., 2019; Yang et al., 2022) fluctuations. Furthermore, it flows at speeds much slower than blood across brain compartments of varying size and shape (Rasmussen et al., 2021), and these slow speeds can be challenging to detect in conventional flow imaging paradigms. Additionally, CSF transport is highly state-dependent (Fultz et al., 2019; Xie et al., 2013), and its dynamics can vary from second to second. Due to the dynamic nature of CSF flow, non-invasive imaging techniques that can image at high spatiotemporal resolution and with high sensitivity are needed to track CSF flow dynamics in humans.

Fast fMRI has recently been shown to be an effective method for detecting CSF flow (Fultz et al., 2019), by leveraging the well-known Time-of-Flight effect (Carr, 2012), where inflow of fresh fluid generates bright signals compared to already saturated stationary tissue. Measuring inflow-weighted signals with fMRI provides high sensitivity and high temporal resolution. Additionally, fMRI-based CSF imaging can measure flow while simultaneously providing whole-brain blood-oxygenation-level-dependent (BOLD) signals. Since vascular dynamics are critical for driving CSF flow, these simultaneous BOLD measures provide rich information about the mechanisms contributing to CSF flow. However, a significant drawback of current fast fMRI flow imaging is that it does not provide quantitative flow measurements. The most common method for quantifying CSF flow with MRI is phase contrast imaging, which has provided insight into CSF flow locked to cardiac or respiratory cycles (Chen et al., 2015; Spijkerman et al., 2019; Sweetman & Linninger, 2011; Takizawa et al., 2017; Yatsushiro et al., 2022; Yildiz et al., 2017), particularly in regions with high velocity flow such as the aqueduct. Phase contrast provides excellent quantitative information but does not simultaneously measure brainwide BOLD signals. Fast fMRI is thus an appealing complementary technique, as its high sensitivity, temporal resolution, and multimodality provides distinct information. Furthermore, since fMRI is widely used and extensive datasets have been collected with BOLD fMRI, both in labs around the world as well as in large consortia such as the Human Connectome Project (Van Essen et al., 2013) and UK Biobank (Sudlow et al., 2015), the ability to extract CSF inflow signals from BOLD data could enable a broad range of new investigations of CSF flow properties in the human brain. Intriguingly, recent studies repurposing public fMRI datasets have indeed found signs of CSF inflow signals (Gonzalez-Castillo et al., 2022; Han et al., 2021), suggesting potential wide applicability of fMRI flow measures that can take advantage of datasets that have already been collected. However, the qualitative nature of the fMRI inflow signal limits the inferences that can be made with current methods.

To accurately interpret CSF flow from fMRI data, it is critical to understand the relationship between fMRI flow-weighted signals (referred to here as ‘inflow signals’) and the true underlying flow dynamics. Inflow signals are not quantitative: they are measured in arbitrary units (Kim et al., 2022) that are not directly proportional to velocity, and the mapping between flow speed and the inflow signal is nonlinear. Important prior work has formulated the relationship between flow velocity and the flow-weighted MRI signal as a function of image acquisition parameters, namely RF pulse history, slice thickness, repetition time (TR), echo time (TE), and flip angle (J. H. Gao et al., 1996; J. H. Gao & Liu, 2012; J.-H. Gao et al., 1988). This prior work considered constant flow velocities, and subsequent work expanding its applications also retained that assumption (Diorio et al., 2024). However, in fast fMRI, finer scale temporal dynamics can be resolved and thus the measured inflow signal can be used to track rapidly varying flow. Additionally, time-dependent flow across slices and multi-slice excitation schemes used in modern fast fMRI paradigms alter the measured inflow signals, requiring an update to how the timing of RF excitations is modeled. This can also have a large effect on the inflow signal caused by bidirectional flows, where fluid flows backwards and exits the imaging volume, leading to much longer longitudinal recovery periods and thus altering inflow signals. Since CSF flow is dynamically modulated, pulsatile, and bidirectional, the ability to interpret and quantify time-varying inflow signals is needed to infer in vivo CSF flow dynamics from fMRI data.

To improve interpretation of dynamic inflow signals in fMRI, we developed a forward model which takes a time-varying fluid velocity as input and simulates fMRI inflow signals. This model is designed to account for simultaneous multislice techniques in modern fMRI acquisitions, to account for dynamic changes in flow and bidirectional flow, and to account for the spatial dependence of velocity on individual anatomy. We validated our model in phantom and in human studies by showing that the model successfully predicts fMRI inflow signals during dynamic, pulsatile flow. Furthermore, we identify features of the signal that are critical to consider when making inferences about flow dynamics in fMRI studies, such as how slice placement within the flow compartment can introduce a volume-dependent bias as well as how differences in signal contributions from slowly vs. rapidly changing velocity can introduce a frequency-dependent bias. To enable flow velocity quantification, we used our forward model to develop a physics-based deep learning framework and demonstrated that it can accurately predict flow velocity directly from fMRI inflow signals. This work enables quantitative assessments of flow velocities using fMRI and informs how we should interpret inflow signals in fMRI datasets, significantly increasing the information that can be gained from neuroimaging studies.

## 2 Methods

### 2.1 Theory

The magnetic resonance (MR) signal comes from detecting how the magnetization of protons within a strong magnetic field respond to radiofrequency (RF) pulses that are repeatedly sent into the imaging volume at time intervals called the repetition time (TR). If repeated RF excitations happen sufficiently close in time, the magnetization from stationary protons in tissue becomes saturated and reaches a steady state, whereby the magnetization remaining at the time of each RF pulse is lower than the initial magnetization at the start of the experiment. Fresh protons that flow into the imaging volume that have not yet experienced any RF pulses will have high signal relative to the stationary protons, and this signal attenuates as protons flow deeper into the imaging volume and accumulate RF pulses. The difference in signal between flowing and stationary protons is the contrast mechanism for the flow-enhancement, i.e., inflow effect (Carr, 2012; Vlaardingerbroek, 2004), which in an fMRI acquisition appears as bright signal at the edge slices of the imaging volume. When protons move quickly through the slices, they accumulate fewer RF pulses within each slice resulting in less saturation and higher signal. When protons move slowly, they spend more time in slices and their signal within those slices become more attenuated. Thus, the magnitude of inflow signal is influenced by the speed of flow. Figure S1 illustrates this concept in the case of constant plug flow. For example, when the flow is slow, fluid within the first slice can be divided into partitions each having different amounts of excitation reducing the overall signal in that slice. With faster flow, fluid can flow through and exit the first slice before the next RF pulse is sent, causing fluid in the first slice to always be excited once which maximizes signal in the first slice. The situation is more complex with dynamic flow, which will be discussed in the next section.

### 2.2 Forward model development

The inflow effect has been known for decades and has been previously modeled in fMRI studies (J. H. Gao et al., 1996; J. H. Gao & Liu, 2012). Prior work assumed that the time in between RF pulses received by the flowing spins is always equal to the TR. However, the time in between excitations received by flowing spins can be variable during a multislice experiment. This becomes particularly important when dealing with bidirectional flows, where fluid can flow backwards outside the imaging volume, allowing for much longer longitudinal recovery periods than TR. Therefore, the inflow signal is affected by the periods of outflow that precede flow into the imaging volume. To account for variable inter-pulse periods, we derived an expression that defines the flow-enhanced MR signal (arising from the transverse magnetization M_T_) as a function of the number of RF pulses received (n) and the time durations in between each of the received RF pulses (Δt):

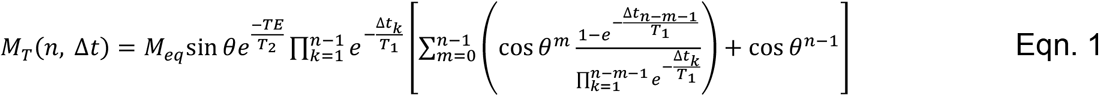

where T_1_ is the longitudinal relaxation time constant, θ is the flip angle, M_eq_ is the equilibrium longitudinal magnetization, TE is the echo time, and T_2_ is the transverse relaxation time constant. The inflow signal is then the difference between M_T_ and the steady state signal from stationary spins (n→∞; Δt = finite, in Eqn. 1). Derivation of Eqn. 1 is described in the Appendix. We also show that the steady state behavior of a previous relationship (Carr, 2012) that assumed all inter-pulse periods to be TR can be recovered by setting Δt = TR (the Appendix). In simulations of CSF flow in this study, we used T_1_ = 4 seconds and T_2_ = 1.5 seconds. For all other parameters in Eqn.1, we used values taken from the fMRI imaging protocol used in validation experiments.

The flow-weighted signal within each imaging slice that is measured in an imaging experiment comes from the combined effect of many spins within the flowing media which can each have different spin excitation histories. Figure 1 illustrates example simulations of flow through a straight tube where a given input velocity can be imposed at every location along the tube. We tested various input velocity waveforms to illustrate how velocity maps to inflow signals across slices. To model the overall signals within imaging slices, we simulate the signal contributions of many individual spin ensembles moving according to an input velocity (Fig. 1 top row). Spin ensembles are represented as point particles that are assigned initial positions behind, within, and ahead of the imaging slices over a distance wide enough so that there are enough of them included for a given simulation duration. Coarse spacing is shown between initialized spin ensembles in Fig. 1 for illustrative purposes. When the simulation begins, each spin ensemble’s position changes from its initial position according to the integral of the input velocity. Each spin ensemble’s position over time is indicated by the black curves in Fig. 1 middle row as they move through the first three imaging slices. The timing of slice-specific RF excitations is given by the imaging protocol and included in the simulation (vertical black lines in Fig. 1 middle row). The profile of the excitations was assumed to be a step function spanning each slice. When a spin ensemble’s position curve intersects an RF excitation, that spin ensemble’s excitation history is updated, and these events are indicated by black dots in Fig. 1 middle row. We track the excitations to define n and Δt in order to repeatedly solve Eqn. 1 for each spin ensemble as they move through the slices over time. The signal evolution of each spin ensemble is illustrated by the transparency of the black curves. The more RF excitations (black dots) accumulated, the more the signal attenuates (more transparency). Substantial signal recovery can be observed during periods of outflow where some spin ensembles near the edge move outside the imaging domain and stop receiving RF excitations. After each repetition cycle, the signals from all spin ensembles are averaged together within each slice, in order to yield timeseries of inflow-weighted signal for each imaging slice (Fig. 1 bottom row). Figure 1c shows that information about flow speed is encoded in the relative signal intensities across imaging slices, i.e. the slice decay rate. However, this slice decay rate as well as the shape of the inflow signal curve is influenced by other factors such as the flow frequency (Fig. 1d) and the directionality (Fig. 1a,b). Additionally, it is intuitive that the fMRI inflow signal only detects signal changes when flow moves into the imaging domain, however Figure 1 shows that the outflow (i.e. negative velocity) does affect signal magnitudes during subsequent inflow. This is due to the signal recovery period that spins near the boundary undergo as they flow below the first imaging slice and thus no longer become excited.

**Figure 1.**
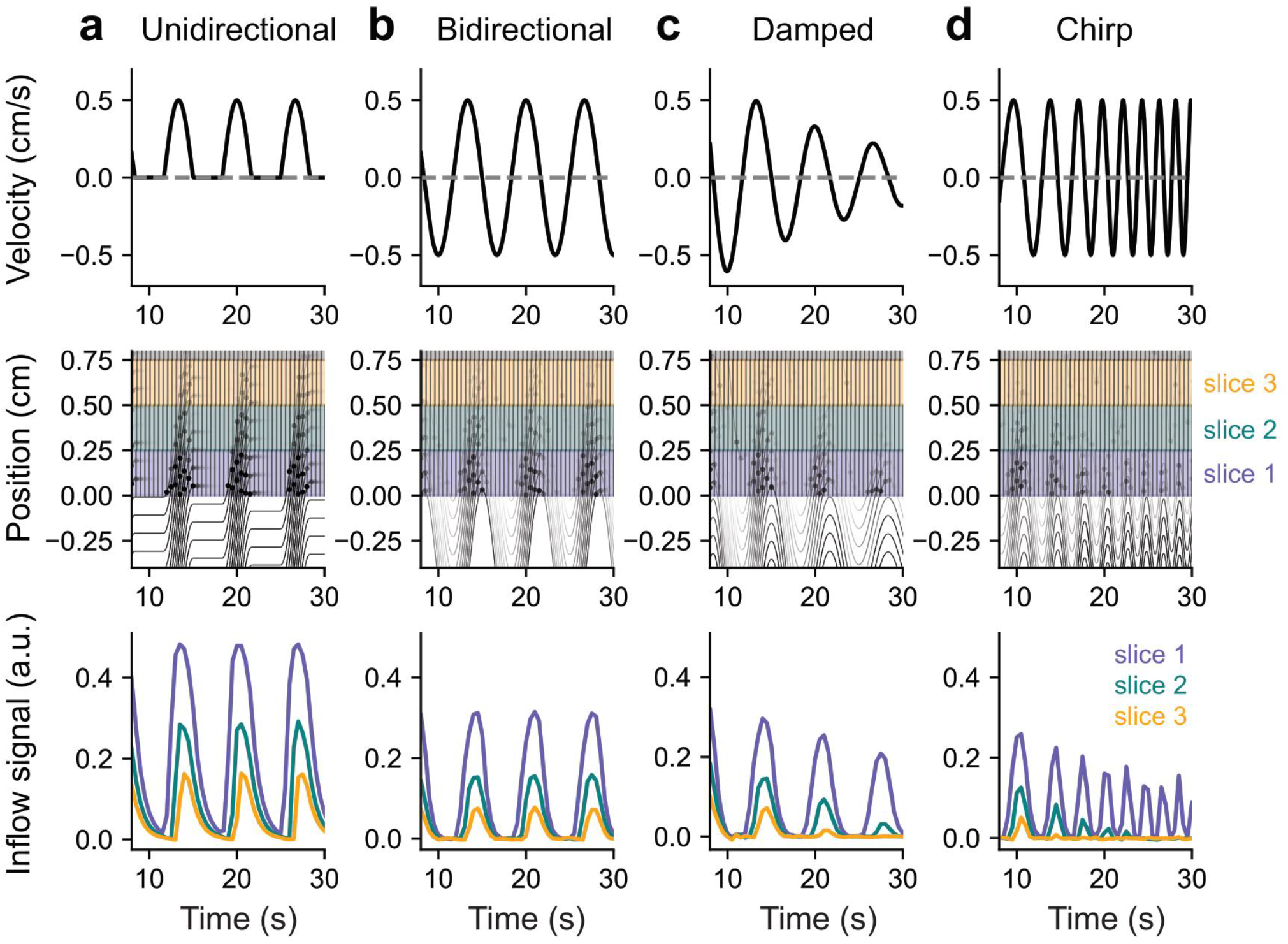
Model schematic showing how an input velocity translates to simulated fMRI inflow signals in the imaging slices. (a-d) Example simulations using unidirectional, bidirectional, velocity amplitude-modulated (damped), and velocity frequency-modulated (chirp) flow inputs into the model. Top row show input velocity waveforms; middle row show spin ensemble position-time curves across the first three imaging slices based on the input velocity; bottom row shows simulated fMRI inflow signals in the first three slices after averaging spin ensemble contributions within each slice over time. Vertical black lines in middle row panels represent slice-specific RF excitation timings; black dots represent each spin ensemble receiving an excitation; the transparency of the position-time curves and associated black dots represents each spin ensemble’s signal evolution. A subset of spin ensembles with coarse spacing are shown here for illustrative purposes. These simulations were performed in a straight tube such that the same input velocity was imposed at every position.

### 2.3 Forward model input velocity

The forward model simulations of flow require integration of a velocity field V(t, x) to define the motion of spin ensembles through imaging slices. This velocity field is a function of time t and position x and is modeled by the following ordinary differential equation:

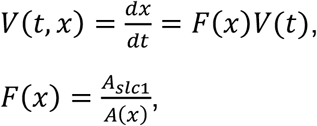

where F(x) is a non-dimensional position-dependent term which depends on the proton’s position (x) along the flow compartment, A(x) is the cross-sectional area as a function of depth extracted from each individual’s anatomical images, and A_slc1_ is the cross-sectional area at the location of the first fMRI slice. By convention, x=0 corresponds to the bottom of the first fMRI slice. This approach accounts for spatial variations in flow velocity due to variable cross-sectional area along the flow compartment. For example, if a spin ensemble flows with velocity V_1_ at position x_1_ with area A_1_, then it should have velocity V_1_/2 at position x_2_ with area 2A_1_. We solved this differential equation using the solve_ivp function from the SciPy Python package (version 1.11.4) where each spin ensemble is given an initial position. We set F(x) equal to 1 for analysis of flow phantom data where flow occurred in straight plastic tubes. During simulation, spin ensembles eventually move well past the imaging slices and reach a position x where there is no available cross-sectional area to use for defining F(x). To ensure that F(x) is defined everywhere, we set the area to 0.05 cm^2^ at positions where there are no cross-sectional areas defined from the anatomical segmentation or if the area was below 0.05 cm^2^. Taken together, the flow of spin ensembles follow the velocity field V(x, t) which depends on the temporal dynamics of the numerical input velocity corresponding to the bottom of the first fMRI slice and position of spin ensembles as they move through the flow compartment.

### 2.4 Inverse model development

We designed a neural network to learn the inverse mapping between fMRI inflow signals and velocities. The input feature vectors of the neural network consist of fMRI inflow signals from the bottom three imaging slices as well as the cross-sectional areas (cm^2^) and corresponding positions (cm) of the areas within the flow compartment (Fig. 6a). The position feature vector for the areas must be shifted such that a position of 0 cm corresponds to the bottom of the first fMRI slice. We set the length of each input feature vector to be 200 which corresponds to 200 time points for the fMRI inflow signals (~100 seconds of data with our TR of 0.504 seconds). The two features of cross-sectional area and corresponding positions were interpolated after extracting from anatomical images to also be of length 200 (see Methods section 2.15 *4th Ventricle cross-sectional area definition*). The output of the neural network is a velocity vector (cm/s) of length 1000 defined to be the same amount of time in seconds as the input.

The neural network consisted of three sequential 1D convolutional layers (kernel size = 3; padding = 1), progressively increasing the number of feature channels. Those layers were then followed by three sequential transpose 1D convolutional layers (kernel size = 3; padding = 1) progressively decreasing the number of feature channels. We performed batch normalization and ReLU activation after each convolution. After the six convolutions, the data was flattened and passed through two fully connected layers before outputting the final velocity timeseries. Input fMRI inflow signals were preprocessed according to Methods section 2.12 *fMRI preprocessing* without lowpass filtering and demeaned before being passed into the neural network. A schematic is shown in Fig. 6a showing the layers of the network along with the dimensions of data as it moves through the network.

We trained the neural network on simulated data generated by the physics-based forward model, which takes an input velocity V_input_, cross-sectional areas, and the positions of those areas shifted such that a position of 0 cm is at the bottom of the first fMRI slice, and predicts the resulting fMRI inflow signal. Each simulated sample uses an input velocity (V_input_) with temporal dynamics defined by a sum of N_f_ sinusoids:

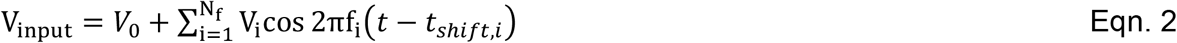

where V_i_ is amplitude, V_0_ is the offset velocity, f_i_ is frequency, t is time, and t_shift,i_ is each sinusoid’s time-shift. We used 100 sinusoids where frequencies ranged between 0.01 and 1 Hz with 0.01 Hz spacing. The V_0_ for each sample was randomly chosen between −0.1 and 0.1 cm/s (based on measured mean phase contrast velocities shown in Fig. S2d), t_shift,i_ (seconds) was randomly chosen between 0 and each sinusoidal period, and V_i_ was randomly sampled from a distribution that will be described in more detail. Rather than sampling the V_i_ values completely randomly which would generate mostly unphysiological training examples, we constrained the sampling based on magnitudes observed in real velocity data measured via phase contrast imaging (Fig. S2a-c). We defined the amplitude sampling bounds such that each f_i_ is randomly assigned a V_i_ between B_L_(f_i_) and B_U_(f_i_), where B_L_(f) and B_U_(f) are frequency-dependent curves defining the lower and upper velocity bounds defined as follows:

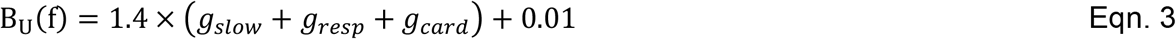

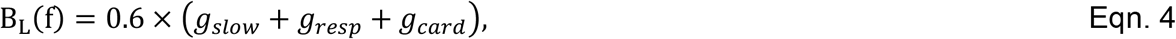

where each g_x_ is a Gaussian distribution corresponding to key dynamics we observed:

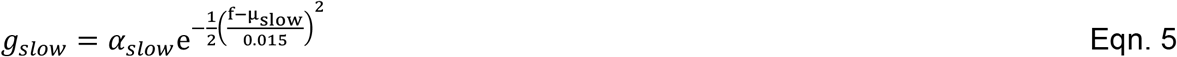

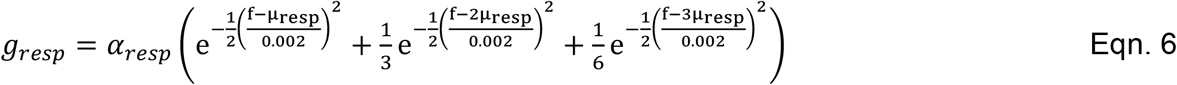

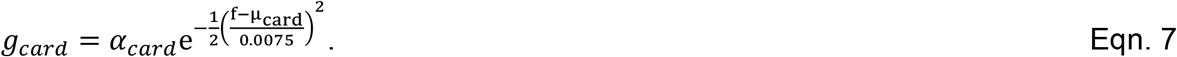

Eqn. 5 represents slow dynamics using a Gaussian distribution centered at µ_slow_, Eqn. 6 represents respiratory dynamics centered at breathing frequency µ_resp_ including harmonics at 2µ_resp_ and 3µ_resp_, and Eqn. 7 accounts for cardiac fluctuations using a Gaussian distribution centered at µ_card_. For each sample, free parameters are chosen using uniform sampling within specific ranges. For the slow distribution: *α*_*slow*_ is sampled between 0 and 0.1 cm/s, and *μ*_slow_ is sampled between 0.035 and 0.065 Hz. For the respiratory distribution: *α*_*resp*_ is sampled between 0 and 1.0 cm/s, and *μ*_resp_ is sampled between 0.1 and 0.3 Hz. For the cardiac distribution: *α*_*card*_ is sampled between 0 and 0.2 cm/s, and *μ*_card_ is sampled between 0.8 and 1.0 Hz. Because most of our human validation data contained flow specifically during 0.167 Hz paced breathing, we also defined a 25% chance that *μ*_resp_ for a given sample is forced to be equal to 0.167 Hz. This approach was not strictly necessary, but was performed here to ensure that enough examples of 0.167 Hz flow were included in the training dataset without having to substantially increase the total number of training samples. Fig. S3 shows one example curve sampled between B_L_ and B_U_, with particular values for the Gaussian parameters.

Each sample also used anatomical information as input into the forward model. To define this anatomical information for each training sample, we used scaled/shifted versions of extracted 4^th^ ventricle anatomy from study participants. For each sample, one of the cross-sectionals area from study participants (Fig. S4) is randomly selected and that area curve is then shifted such that the location corresponding to the bottom of the 1^st^ fMRI slice can vary between −1 and +1 cm from the widest location (an interval that covers actual slice placements we used in the study). The area curve is then multiplied by a factor that varied between 0.8 and 1.2. The approach described thus far was applied to training samples meant to approximate human data, however because we also applied the same network to flow phantom data which has distinct differences from human data, we created a subset of samples tailored to phantom data. During sampling, we set a 25% chance of being in “ phantom mode” where rather than multiple gaussians, we only used a single gaussian defined as follows:

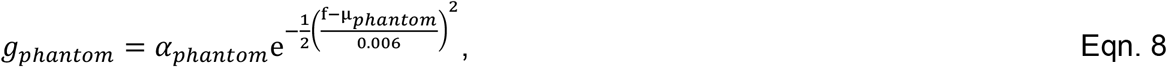

where *μ*_*phantom*_ was randomly chosen to be either 0.05, 0.1 or 0.2 Hz, and *α*_*phantom*_ randomly varied between 0 and 1.2 cm/s. Also, in “ phantom mode” the V_0_ was forced to be 0 because the flow phantom pump produced purely bidirectional oscillations, and the anatomical information for phantom samples was set to be a straight tube. Finally, for both human and phantom samples, we added zero-mean normally distributed gaussian noise to the simulated inflow signals (max amplitude was 1 for human samples and 0.25 for phantom samples; standard deviation varied between 0.01 and 0.1). We trained the neural network on 45,000 samples using PyTorch (version 2.1.2) with batch size = 16, number of epochs = 120, learning rate = 0.001, and using the Adam optimizer. We separated 10% of the simulated dataset as test data and used the mean-squared error (MSE) loss criterion. The MSE of model output evaluated using the test dataset was 0.016 cm^2^, indicating good performance of the neural network on the simulated test data. We also confirmed network convergence by inspecting the loss curves after training.

### 2.5 Phantom data acquisition

We performed validation experiments on a flow phantom to assess model accuracy and performance. The flow phantom system consisted of a controllable syringe pump (Harvard Apparatus - Pump 11 Elite Infusion/Withdrawal Programmable Single Syringe) connected to a plastic tube that was routed into the scanner. The tubing in the bore was fixed onto a gel phantom such that tubing came into the bore and then wrapped back around to the control room. Our EPI slices were then positioned to intersect both the “ in” and “ out” portions of the tubing. The fluid used was tap water. To produce oscillatory-like flows using the pump, we defined an infuse and withdraw cycle on the system’s interface. We adjusted the volume flow rate and the time period of the infuse and withdraw phases of the cycle to control the velocity and frequency of flow, respectively. Velocities tested ranged between 0.1 to 1.6 cm/s; frequencies tested were 0.05, 0.1, and 0.2 Hz. For cases with constant velocity, the system was set to infuse at a specified flow rate for a set period of time. We acquired a total of 40 flow conditions (30 oscillatory flow; 10 constant flow). The experiments were performed on a 3T Siemens Prisma scanner with a 64-channel head coil.

### 2.6 Human MRI acquisition

Eight healthy adult human participants (4 male, 4 female, ages 21 - 42) provided informed written consent and all experimental procedures were approved by the Boston University and Massachusetts General Hospital Institutional Review Board. Experiments were performed on a 3T Siemens Prisma scanner with a 64-channel head coil. Anatomical images were acquired with a 1 mm isotropic multi-echo MPRAGE (MEMPRAGE) anatomical scan (van der Kouwe et al., 2008). After visual identification of the fourth ventricle, subsequent flow measurement runs were manually positioned to intersect orthogonally with the ventricle direction of flow. fMRI inflow signals were measured using a gradient-echo EPI sequence (TR=0.504s, TE=30ms, 2.5mm isotropic voxels, 21 slices, FA=45, matrix=92×92, MB=3 with FOV shift factor=4). Phase contrast runs used a TR=19.8ms, 2.5mm isotropic voxels, single-slice, matrix=68×80, VENC=6cm/s. We acquired phase contrast images using a Siemens BEAT sequence with a standard multi-shot TrueFISP readout (echo time = 7.1ms). Phase encoding lines were acquired sequentially from bottom to top of k-space, with the central lines captured mid-acquisition. The time for acquiring both scans with opposing gradient pairs (i.e. temporal resolution) was 1.0098s.

### 2.7 Experimental protocol

We acquired data during runs of a paced breathing task and separate runs of rest. In the paced breath task, to drive CSF pulsations with controlled timing, the subject performed 6 minutes of paced breathing with 6-second breath cycles (3-second breath in; 3-second breath out) (Dreha-Kulaczewski et al., 2017). Breathing task instructions were displayed on a screen that the subject could view during the scan. The resting state runs lasted 6 minutes, and the participants were instructed to rest with their eyes open. For each condition of either paced breathing or rest, we acquired a single run of phase contrast imaging and then immediately afterwards a single run of fMRI. Velocities from phase contrast imaging were treated as ground truth measures of velocity associated with each of the subsequent fMRI runs. Importantly, the phase contrast and fMRI are separately acquired runs: they allow measurement of average flow dynamics and speed for a given task (breathing or rest), individual participant, and slice position, but not cycle-to-cycle comparisons. Physiological recordings were simultaneously recorded during fMRI acquisitions. Respiration was measured through a piezoelectric belt (UFI systems, Morro Bay, CA, USA) around the abdomen. Pulse was measured with a piezoelectric pulse transducer around the left thumb (AD Instruments, Colorado Springs, CO, USA). Both physiological recordings were acquired using LabChart 8.0. Respiratory recordings were unavailable for two of the subjects due to equipment malfunction.

### 2.8 Anatomical image preprocessing

Anatomical MEMPRAGE images from human sessions were bias-corrected using SPM12 in MATLAB and then automatically segmented using Freesurfer (Fischl, 2012).

### 2.9 Slice positioning

Flow properties of CSF can vary with respect to location in the brain and therefore measured inflow signals can vary substantially based on where the volume is positioned. In our human experiments, we positioned edge slices at the 4th ventricle where CSF flow has been previously measured (Chen et al., 2015; Fultz et al., 2019). The cross-sectional area within the fourth ventricle also varies with depth, therefore variations in slice-placement along the fourth ventricle can result in differences in flow velocity: wider regions have slower flow, while narrower regions have faster flow. This effect can confound comparisons between individuals and was therefore accounted for in this study. For every participant, we manually positioned the edge slice to intersect a relatively narrow location in the fourth ventricle so that we could measure higher velocity flow and increase the amplitude of measured inflow signals. For five of the eight human participants, in addition to the acquired data at the narrow slice placement, we also acquired data near the widest part of the fourth ventricle to directly assess flow differences caused by differences in slice placement. All slice positioning was manually performed while referring to anatomical images acquired at the beginning of each scan. We first positioned the fMRI slices, and then set the single phase contrast imaging slice positioning to match the first fMRI slice. Fig. S4 shows every participant’s area-depth curve with narrow and wide (where applicable) slice placements indicated.

### 2.10 Flow ROI selection

To extract CSF flow signals from the fMRI and phase contrast imaging data, we first defined a flow ROI in the fMRI data by manually selecting the brightest voxels within each of the first three imaging slices. Voxels were selected conservatively to minimize partial volume effects, keeping only the most CSF-filled voxels with highest flow signal (5.41 ± 3.45 voxels on average per slice). We next defined the flow ROI in the single-slice phase contrast imaging data by registering the first slice of the fMRI flow ROI mask to the phase contrast slice using ANTs (version 2.3.5). The registered phase contrast ROI mask was then manually adjusted to ensure that the flow voxels spatially matched those selected in the first fMRI slice, since we directly compare the two signals in our analysis. In addition to the flow voxels, we also labeled stationary voxels nearby the flow ROI for phase contrast imaging offset correction.

### 2.11 Partial volume correction

To correct for partial volume effects from tissue in the fMRI data, we implemented an anatomy-based algorithm that takes advantage of the difference in signal between CSF and tissue. We did not implement this algorithm for the phase contrast data because it was difficult to accurately register the anatomical image to the single slice in phase contrast imaging. To mitigate partial voluming in the phase contrast data, we instead relied on the conservative ROI selection. An initial registration matrix between the functional and T1-weighted MEMPRAGE image was calculated using boundary-based registration. We then resampled the anatomical image into functional space, where we defined two reference T1-weighted values: the first from a tissue voxel near the flow ROI (T1W_,tiss_), and the second being the minimum T1-weighted value within all manually labeled CSF voxels (T1W_,CSF_) which was assumed to be the most CSF-filled voxel (since CSF is dark on MEMPRAGE images). A scaling factor was defined based on a linear interpolation between the tissue and CSF reference values:

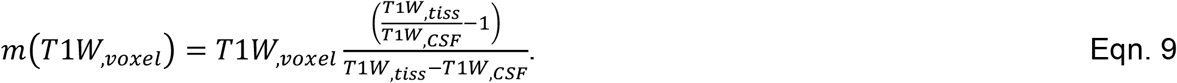

The fMRI timeseries of each voxel in the CSF flow ROI was then multiplied by m with T1W_,voxel_ being the T1-weighted value for the voxel. This procedure scales signals up the more partial voluming is present (i.e., partial voluming is assumed to increase as T1W_,voxel_ approaches T1W_,tiss_).

### 2.12 fMRI preprocessing

After correcting for partial voluming, fMRI signals from the flow ROI were averaged slice-wise to yield one timeseries for each slice (the bottom 3 slices). The first 40 datapoints of each fMRI timeseries were removed to omit the initial transient period of high signal. We then computed the mean value from the 10% lowest amplitude values of each slice’s timeseries and then respectively for each slice we subtracted this mean to establish a baseline near zero. The resulting inflow signals were lowpass filtered (5^th^ order Butterworth filter) below 0.5 Hz to stay below the Nyquist frequency of the phase contrast imaging data. Runs acquired at the same slice positioning with the same task were concatenated. Motion statistics for paced breathing runs in human sessions were obtained using AFNI version 19.1.00 for calculation of frame-wise displacement, and high-motion epochs were manually removed from the fMRI data. We opted to not use motion-corrected data in the analysis because motion correction is prone to being inaccurate at edge slices where tissue moves in and out of the imaging volume, causing nonflow related signal intensity changes. We did not perform slice-time correction because slice-specific RF excitations are accounted for in simulations by taking the RF slice timings as an input parameter.

### 2.13 Phase contrast image preprocessing

The phase signals from voxels in the flow ROI were averaged together to obtain a phase timeseries for the single slice. A stationary offset phase signal was obtained by averaging the phase signals in the labeled stationary voxels nearby the flow ROI, and was then subtracted from the phase timeseries extracted from the flow ROI. The resulting offset-corrected phase signal was then scaled by VENC to convert to units of velocity. Runs acquired at the same slice positioning with the same task were concatenated. We did not perform any filtering on phase contrast velocity data.

### 2.14 Cycle averaging

To obtained cycle-locked average phase contrast velocity and fMRI inflow signals from the oscillatory data (either paced breathing runs in human sessions, or oscillatory flow phantom runs), we first upsampled the timeseries by a factor of 4 using linear interpolation. Next, we identified peaks in the signal using the find_peaks function in the SciPy Python package (version 1.11.3) and defined each oscillatory cycle as the data points between identified peaks of the signal in the first slice. The optional distance parameter in the find_peaks function was set to be 1 second less than the prescribed controlled timing of the oscillations (i.e., a 5 second threshold for paced breathing data where the instructed breathing cycle was 6 seconds). This distance parameter defines the minimum time period between neighboring peaks. For example, in human paced breathing data, this distance threshold ensured that short breathing periods resulting from poor task compliance were not included in the average oscillation. The prominence parameter in find_peaks was set to 0.1 for both phase contrast velocities (in cm/s) and the normalized inflow signals. All the oscillation periods were then averaged together to obtain a mean cycle-locked average signal. Power spectral densities were computed using the welch function from the SciPy Python package (version 1.11.3). Respiratory belt signals were missing from two subjects scanned due to equipment malfunction. For this reason, to avoid excluding data, we opted to use this peak detection approach for computing period averaged signals for consistency across all subjects.

### 2.15 4^th^ Ventricle cross-sectional area definition

Cross-sectional area of the 4^th^ ventricle as a function of depth was extracted using the segmentation output from Freesurfer’s recon-all. Anatomical voxels labeled as 4^th^ ventricle from the segmentation volume were extracted. To correct for partial volume effects from tissue, we implemented an anatomy-based algorithm that takes advantage of the difference in MEMPRAGE intensity (van der Kouwe et al., 2008) between CSF and tissue. For each slice of the extracted 4^th^ ventricle volume, the minimum T1-weighted value was taken as a reference (T1W_,ref_) and each voxel’s area in that slice was scaled by T1W_,ref_/TW1_,voxel_. This scales down cross-sectional area contributions from 4^th^ ventricle voxels that contain some tissue. The scaled area contributions for each slice of the extracted 4^th^ ventricle volume were summed slicewise to obtain cross-sectional as a function of depth.

## 3 Results

### 3.1 Model validation in a flow phantom during constant and oscillatory flow

We first performed validation experiments on a flow phantom to assess model accuracy, testing whether we could accurately predict fMRI inflow signals for a wide range of input velocity profiles that we controlled. To evaluate how the model performs in the simplest case, when the velocity is constant, we first set the flow phantom to deliver fluid with a single velocity (0.6 cm/s). We analyzed the resulting inflow signals within the flow ROI (Fig. 2a) and further validated this input velocity using phase contrast imaging (Fig. 2b). We then simulated the inflow signals for the bottom three slices of the acquisition volume, using the measured velocity from the phase contrast scans as input (Fig. 2c). The simulated fMRI inflow signal timeseries in each slice was normalized to values in the first slice, and was compared to measured fMRI inflow signals normalized in the same way (Fig. 2d). We found close agreement between our model predictions and measured fMRI data when the flow velocity was constant (Fig. 2b-d, Fig. S5), as expected based on prior work establishing the basis of the fMRI inflow signal (J. H. Gao et al., 1996).

**Figure 2.**
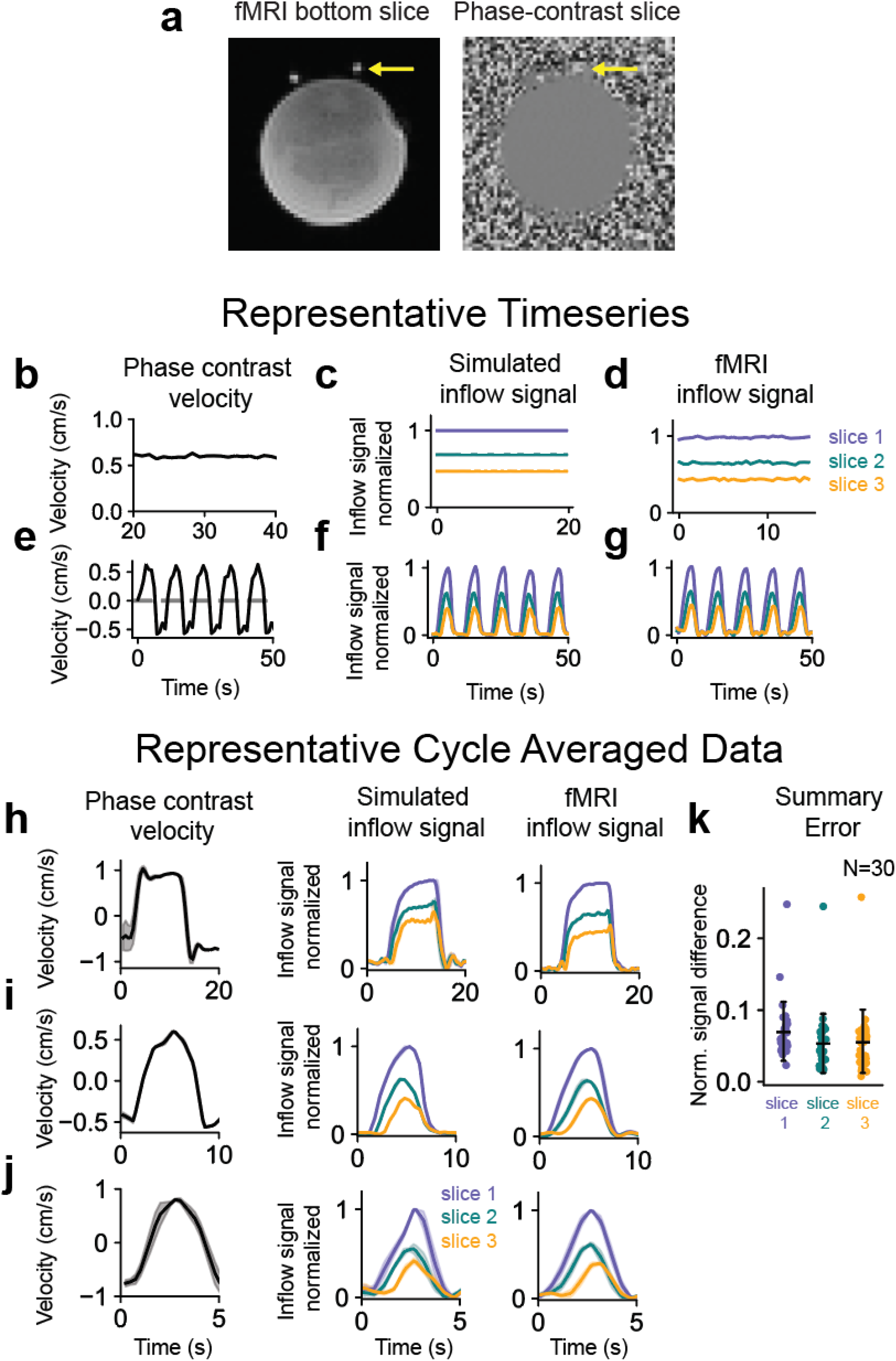
Model validation in a flow phantom during constant and oscillatory flow. (a) Example fMRI and phase contrast images acquired during flow phantom experiments using a syringe pump with flow ROIs in plastic tubes indicated by yellow arrows. (b) Measured phase contrast velocity during constant flow. (c) Simulated inflow signal using the average velocity from panel b (0.6 cm/s) as model input. (d) Measured fMRI inflow signal during the same constant flow condition used in panel b. (e) Measured phase contrast velocity during oscillatory flow from a representative run. (f) Simulated inflow signal using the oscillatory velocity timeseries from panel e as model input. (g) Measured fMRI inflow signal during the same oscillatory flow condition used in panel e. (h-j) Representative cycle averages for the 0.05 (h), 0.1 (i), and 0.2 Hz (j) flow conditions. All simulated and measured fMRI inflow signals are normalized to the mean value of the top 5% of signals in the first slice. (k) Summary errors for flow phantom data in three slices between measured and simulated fMRI inflow signals, using measured velocities as model input (30 oscillatory flow conditions consisting of 10 runs each of 0.05, 0.1, and 0.2 Hz). Mean errors for the three slices were 0.0703 ± 0.0404, 0.0536 ± 0.0407, and 0.0564 ± 0.0433, respectively. Reported errors were computed by taking the mean absolute difference between simulated and measured normalized cycle-averaged traces for each slice.

We next tested time-varying flow in the phantom, which is more representative of the pulsatile flow dynamics observed in humans. Model input was a velocity timeseries obtained from phase contrast data, with velocity set to oscillate at 0.1 Hz (Fig. 2e), and the resulting simulated inflow signals (Fig. 2f) showed close agreement in its dynamics and amplitudes to fMRI inflow signals separately measured during the same flow condition (Fig. 2g). To quantify overall model performance during oscillatory flow (30 runs using either 0.05, 0.1, or 0.2 Hz oscillations), we defined an error metric as the mean absolute difference between the measured and simulated cycle averaged inflow signals (representative examples shown in Fig. 2h-j), and plotted the resulting errors across three slices for all flow conditions (Fig. 2k). These error distributions from phantom data showed that the model was able to accurately capture the relative fMRI inflow signal intensities during dynamic flow. Raw traces from measured and simulated data also qualitatively illustrate good model performance across a wide range of velocities and frequencies (Fig. S6).

### 3.2 Model predicts fMRI flow signals in humans during paced breathing

We next tested whether the model could successfully predict human imaging data, where CSF flow is always pulsatile rather than constant, and where the flow compartment has an irregular shape in contrast to the cylindrical tubes used for phantom experiments. We positioned the imaging slices at the 4th ventricle where CSF flow has been previously measured (Chen et al., 2015; Fultz et al., 2019). To drive CSF pulsations with known timing, we instructed the participants to perform a paced breathing task (Fig. 3a), since CSF flow is known to be modulated by respiration (Chen et al., 2015; Dreha-Kulaczewski et al., 2015, 2017; Kao et al., 2008; Spijkerman et al., 2019). The phase contrast signals taken from an ROI in the 4^th^ ventricle (Fig. 3b) and respiratory belt recordings confirmed a robust induction of CSF oscillations locked to the respiratory task (Fig. 3c,d). To validate model performance, we again used the phase contrast velocity timeseries as inputs into the forward model. Importantly, since phase contrast and fMRI data were acquired in separate runs, and human CSF flow is variable from run to run, we focused on the mean breath-locked CSF flow to calculate performance of the model. We found that we could successfully predict fMRI inflow signals during this task, with close agreement between the model output and the measured signals (Fig. 3e,f). To quantify overall performance across the 8 individual participants at both the narrow and wide slice placements (8 narrow ROIs; 5 wide ROIs), we used the same error metric used in the flow phantom analysis, and found accurate prediction of the fMRI inflow signal (Fig. 3g,h). Summary errors for human data were found to be higher than in phantom data, and mean cycle averaged simulations exhibited a slightly different shape than measured data. These differences could be partially due to the assumption of laminar flow through the imaging slices, which was more valid for flow phantom experiments but less valid in the 4^th^ ventricle where there are much more complex flow patterns. We also note that since the phase contrast and fMRI runs were acquired separately, our error calculations represent an overestimate of the error, since the true velocity varies slightly across runs. Cycle averaged data used to compute summary errors are shown in Fig. S7.

**Figure 3.**
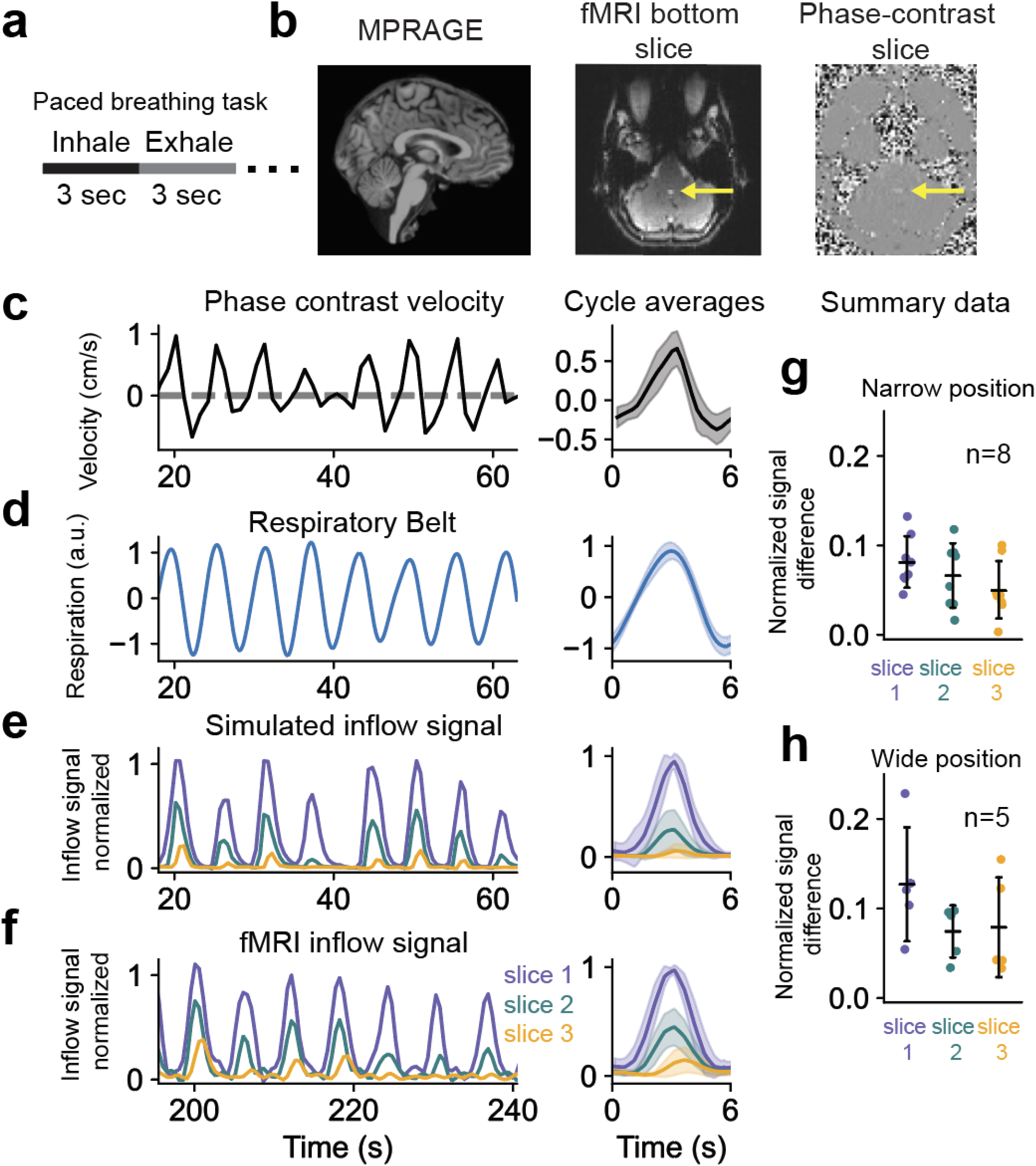
Model predicts pulsatile fMRI inflow signals in humans during paced breathing. (a) Illustration of the 6-second paced breathing task (3 seconds breath in; 3 seconds breath out) presented to human participants. (b) Example brain-masked and intensity-normalized anatomical MEMPRAGE image, fMRI, and phase contrast images from one subject, with flow ROIs at the 4^th^ ventricle indicated by yellow arrows. (c) Measured phase contrast velocity during paced breathing from a representative human participant. (d) Respiratory belt recording during paced breathing from a representative human participant. (e) Simulated inflow signal in the first three imaging slices using the velocity from panel c as model input. (f) Measured fMRI inflow signal during the paced breathing task (acquired in a separate run). Cycle averaged data for each timeseries is shown on the right side of panels c-f. All simulated and measured fMRI inflow signals are normalized to the mean value of the top 5% of signals in the first slice. (g, h) Errors from 8 runs with the edge slice positioned at a relatively narrow position in the 4^th^ ventricle (g) and from 5 runs at a wider positioning. Errors are taken from signal across three slices between measured and simulated fMRI inflow signals during paced breathing that used measured velocities during paced breathing as model input. Mean errors with the narrow slice positioning for slices one, two, and three were 0.082 ± 0.027, 0.066 ± 0.034, and 0.050 ± 0.030, respectively. Mean errors with the wide slice positioning for slices one, two, and three were 0.13 ± 0.057, 0.074 ± 0.026, and 0.079 ± 0.050, respectively. Reported errors were computed by taking the mean absolute difference between simulated and measured normalized cycle-averaged traces for each slice.

### 3.3 Effect of anatomy on fMRI inflow signals

Having established a model that can link specific flow velocities to fMRI inflow signals, we next examined how individual differences in the anatomy of the flow compartment can affect measured fMRI inflow signals. Importantly, while an increasing number of studies aim to compare fMRI inflow signals across groups of patients, differences in anatomy and slice position across subjects are common and can substantially alter measures of inflow signal amplitudes. We focused on image position and anatomical variations in the size and shape of the 4^th^ ventricle, which have important consequences for interpreting inflow signals: a difference in inflow signals across individuals could be due to either a true different CSF flow velocity profile, or simply due to different ventricle size or different image position relative to the ventricle. These variations result in different cross-sectional areas that affect flow velocity: wider regions have slower flow, while narrower regions have faster flow. Furthermore, if the flow compartment widens or narrows relative to the fMRI edge slice, this can also affect the relative signals across slices due to spins speeding up or slowing down as they flow through the slices.

To understand how the spatial dependence of velocity affects measured fMRI inflow signals, we acquired data from two different slice positions, one near the widest part of the 4^th^ ventricle, and another at a narrower part (Fig. 4a). We expected to see faster velocities in the narrower regions due to the increase in flow resistance (Fig. 4b), and indeed, Fig. 4c confirms that flow velocities were higher in data from narrower slice positions than wider ones (compare dots to squares) and that, as expected from mass conservation, multiplying the velocities (Fig. 4c) by corresponding cross-sectional areas (Fig. 4e) yielded volumetric flow rates (Fig. 4f) that were similar between the narrow and wide slice positioning. However, multiplying fMRI inflow signals (Fig. 4d) by cross-sectional areas did not yield results consistent with mass conservation (Fig. 4g).

**Figure 4.**
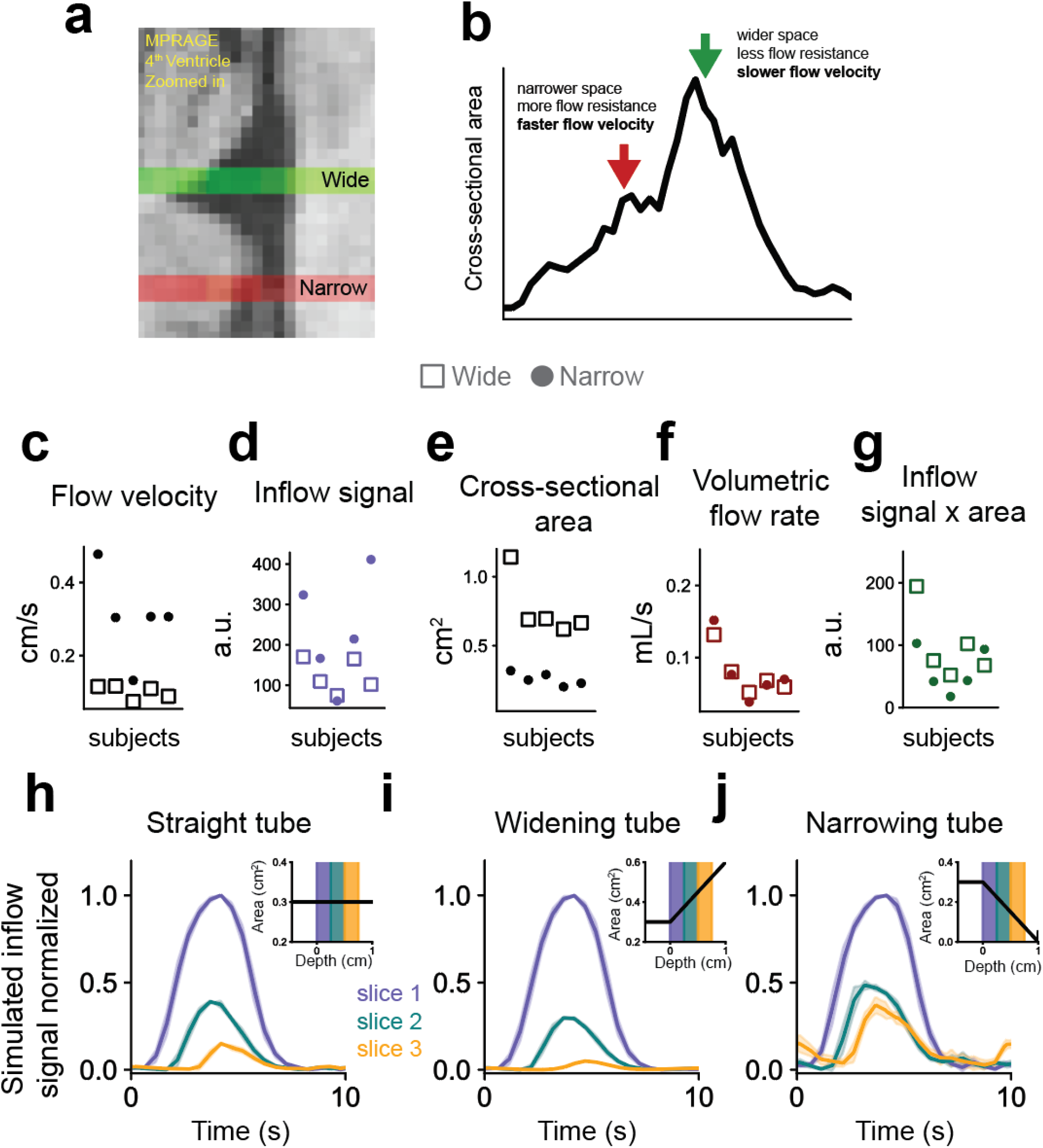
Effect of 4^th^ ventricle size on measured flow velocity. (a) Example from an MEMPRAGE image showing wide and narrow slice positioning in the 4^th^ ventricle used in human experiments. (b) Cross-sectional area vs. depth along 4^th^ ventricle illustrating the differences in flow properties between the wide and narrow slice positioning. (c) Average peak velocity from phase contrast measurements at the wide (squares) and narrow (dots) slice positions, for five subjects during paced breathing. (d) Average peak fMRI inflow signal from the first slice at the wide (squares) and narrow (dots) slice positions, for five subjects during paced breathing. (e) Cross-sectional areas at the wide and narrow positions. (f) Volumetric flow rates at the wide and narrow positions, computed by multiplying panel c with e. Measured velocities were higher at the narrow position than the wide while cross-sectional areas were lower, and volumetric flow rates were similar between the two positions. (g) fMRI inflow signal amplitudes scaled by cross-sectional areas, computed by multiplying panel d with e. (h-j) Cycle-averaged simulated inflow signals using either a straight tube (panel h), widening tube (panel i), or a narrowing tube (panel j) as the flow compartment. Insets illustrate the cross-sectional area with depth of each flow compartment used in simulations, with shaded area indicating the imaging slices.

Importantly, the spatial dependence of inflow signals could create a significant confound in fMRI studies if not accounted for, because inflow signals vary substantially across slices due to flowing fluid speeding up or slowing down as it moves through a space with varying cross-sectional area. This effect was therefore incorporated in the forward model (see Methods) and was used in the simulations with human data (Fig. 3). To investigate the effect of spatial-dependence on the fMRI inflow signal, we simulated inflow signals in three different idealized flow compartments: a straight tube, a widening tube, and a narrowing tube (Fig. 4h-j). We found that when the flow compartment widened relative to the edge slice, there was increased signal decay across imaging slices (Fig. 4i), due to spins slowing down as the cross-sectional area increased. Conversely, a narrowing tube resulted in less signal decay across slices (Fig. 4j). Anatomical variation and positioning should therefore be considered when comparing fMRI inflow signals across subjects, or modeled explicitly using this framework. Individual anatomy is particularly important to consider because the information about flow velocity lies in the relative inflow signal intensities between adjacent slices, which is strongly modulated by where slices are positioned.

### 3.4 Frequency-dependent attenuation in flow-enhanced fMRI signal amplitude

An advantage of the fMRI-based inflow measurement is its high temporal resolution, enabling study of rapid, pulsatile flow dynamics. However, the mapping between oscillating velocity changes and fMRI flow signals has not yet been established. CSF flow can pulse at a wide range of different timescales, including rapid cardiac fluctuations (~1 Hz) (Iliff et al., 2013; Mestre et al., 2018) and slow hemodynamic oscillations (~0.05 Hz) (Fultz et al., 2019; Holstein-Rønsbo et al., 2023; Van Veluw et al., 2020; Williams et al., 2023; Yang et al., 2022). We simulated how fMRI inflow signals would change when flow oscillates at high or low frequencies, and observed a relationship between the oscillatory frequency of an input velocity and the corresponding simulated inflow signal (Fig. 5a). This simulation took as input a velocity defined as a sum of six sinusoids with equal amplitudes over a range of frequencies, and showed that the resulting simulated inflow signal exhibited a frequency-dependent attenuation (Fig. 5a). Specifically, fMRI signals showed a dropoff at high frequencies, such that higher-frequency oscillations produced relatively smaller inflow signals. This simulation suggested that fMRI inflow signals represent a weakly low-pass filtered version of the underlying fluid flow dynamics.

**Figure 5.**
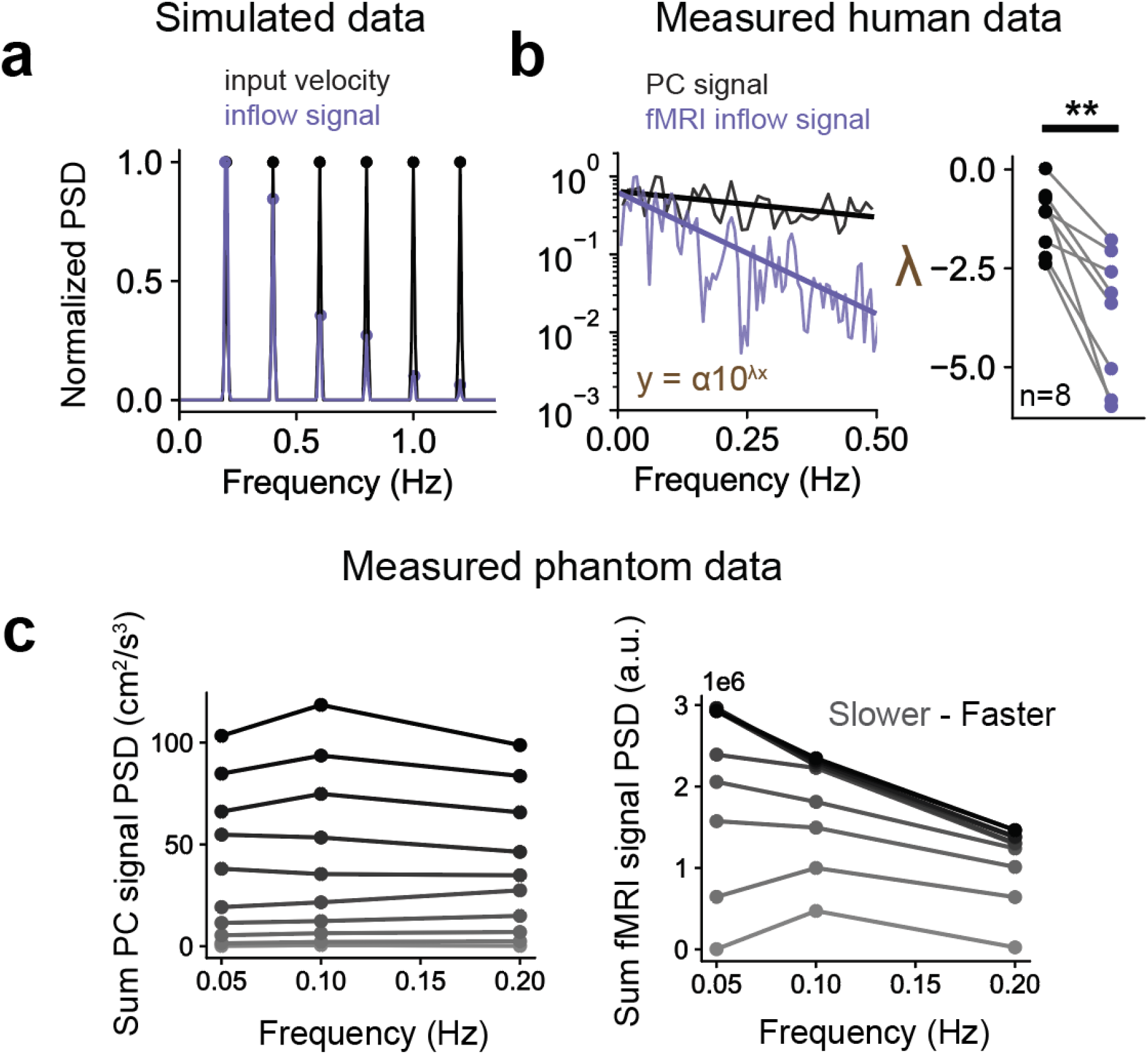
The fMRI inflow signal has attenuated amplitudes when fluid velocity oscillates at high frequencies. (a) Normalized power spectral density (PSD) of a simulated input velocity (black) defined as a sum of six sinusoids oscillating at 0.2, 0.4, 0.6, 0.8, 1.0, and 1.2 Hz, with equal power at each frequency; normalized PSD of simulated fMRI inflow signal in the first slice (purple) using the input velocity (black) as model input. (b) Left panel: Log-scale PSD of a representative phase contrast (PC) velocity (black) and fMRI inflow signal (purple) both measured during eyes open resting state, and each normalized to the maximum power, with power law fits of each spectrum overlayed as colored lines; right panel: group results (8 runs) showing gamma exponents from all the power law fits of phase contrast velocity data (black dots) and fMRI inflow signals (purple dots). Gamma exponents from fits of fMRI inflow signal spectra were significantly lower than phase contrast (p=0.0018; t=4.8883; paired t-test; n=8). (c) Left panel: sum PSD phase contrast velocities from flow phantom experiments at 10 different flow velocities across 3 flow frequencies. (c) Right panel: sum PSD fMRI inflow signal from the first slice from flow phantom experiments at 10 different flow velocities across 3 flow frequencies. The more gray the curve, the slower the flow velocity. The more black the curve, the faster the flow velocity. Summed power spectra values show that fMRI inflow signals were attenuated at the higher flow frequency, despite the speed of input flow being the same.

To validate this relationship in human data, we first examined spontaneous CSF dynamics in the awake resting state. For each participant, we fit a power law to the normalized power spectral density of phase contrast velocities and measured fMRI inflow signals to test whether the frequency content was different between them (representative example in left panel of Fig. 5b). We found stronger attenuation in power of fMRI inflow signals at higher frequencies; pair-wise comparison of mean power law exponents from each subject showed that fMRI inflow signals exhibited significantly (p=0.0018; t=4.8883; paired t-test; n=8) stronger frequency attenuation than velocity (right panel of Fig. 5b).

We also confirmed this frequency attenuation effect in flow phantom data where we could control the flow inputs. We analyzed the summed power spectral density of raw fMRI inflow signals measured across a range of frequencies and velocities. Based on the frequency attenuation we previously observed in human data (Fig. 5b) and simulations (Fig. 5a), we expected to see less raw fMRI signal in the phantom runs using the same input flow speed but higher oscillation frequencies. Indeed, the raw power of fMRI inflow signal was markedly reduced across flow frequencies (0.05, 0.1, and 0.2 Hz) for each flow velocity, while the velocity power was confirmed to be stable across frequencies (Fig. 5c). We also note that the overlapping lines in the right panel of Fig. 5c are due to velocities reaching the critical velocity and thus the inflow signals saturated. Taken together, both human and phantom data corroborate the frequency attenuation effect seen in model simulations. These results demonstrate that a quantitative framework is needed to interpret relative signal intensities in fMRI flow datasets subject to this frequency-dependent bias.

### 3.5 Predicting flow velocity from fMRI inflow signals using physics-informed deep learning

Having shown that our forward model can accurately simulate inflow signals (Fig. 2; Fig. 3), and given that we identified volume-dependent (Fig. 4) and frequency-dependent (Fig. 5) properties of the fMRI inflow signal which make direct assessment of the underlying fluid flow difficult, we next aimed to develop an inverse procedure that can estimate velocity given fMRI inflow signals and individual anatomy. We developed a physics-based neural network framework that takes fMRI inflow signals across three imaging slices in addition to anatomical information as input to predict a flow velocity timeseries (Fig. 6a). We trained the network using a labeled training dataset consisting of simulated data generated using our forward model, allowing us to generate a large dataset for training (45,000 simulated samples).

**Figure 6.**
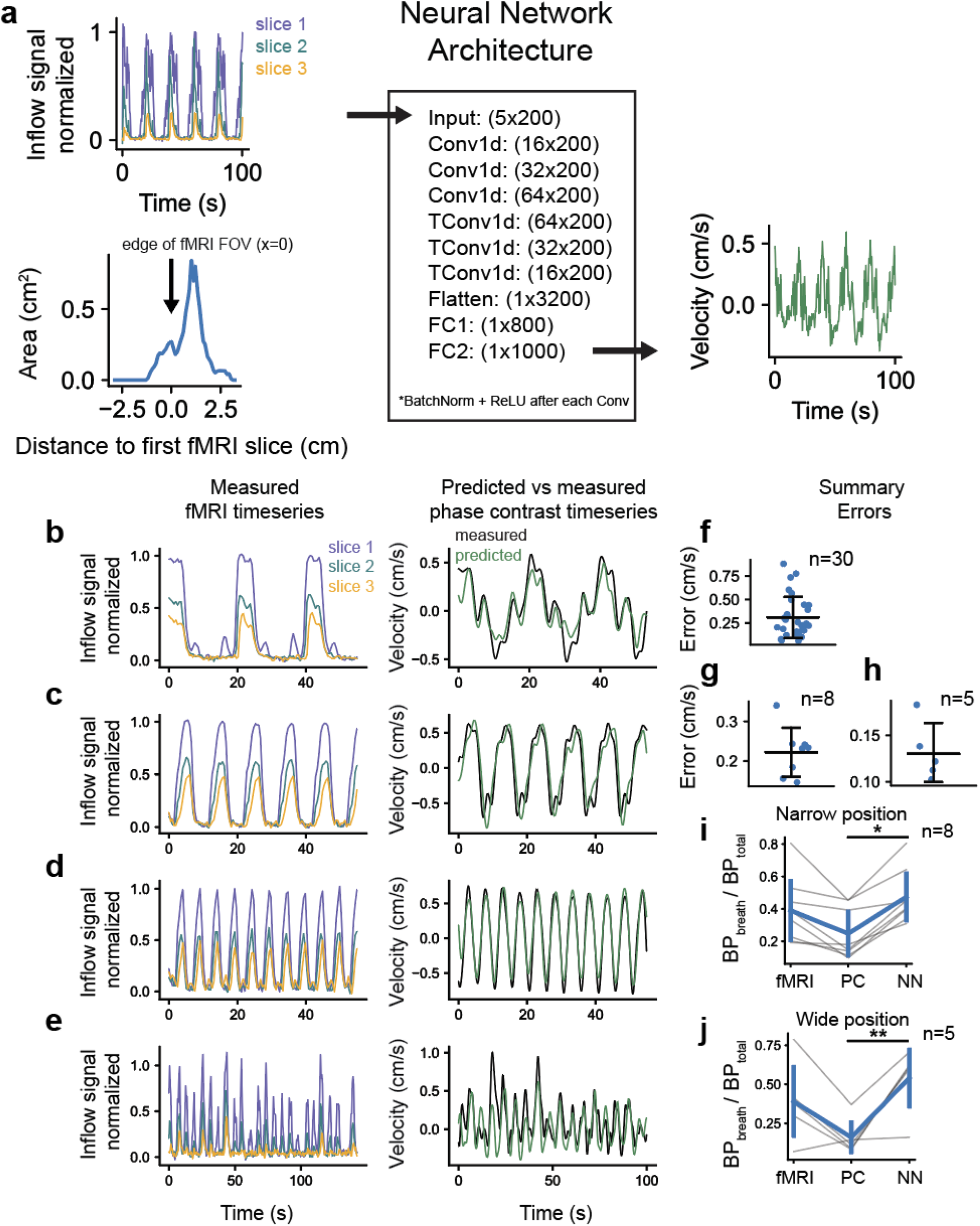
The neural network accurately predicts CSF velocity from fMRI data. (a) Schematic showing the layers of the neural network as well as an example input and output (Conv1d = 1D convolution; TConv1d = 1D transposed convolution; FC = fully connected layer). There are five input channels of length 200 consisting of three fMRI inflow signal timeseries from three consecutive slices (top left panel) as well as cross-sectional areas and corresponding distances from the first fMRI edge slice (bottom left panel), and one output channel of length 1000 yielding a velocity timeseries in cm/s (right panel). (b-e) Left panels: Representative measured fMRI inflow signals from three slices; Right panels: Predicted velocity using measured fMRI inflow signals as input into the neural network (green) and measured phase contrast velocity (black). Panels b, c, and d show flow phantom data using 0.05, 0.1, and 0.2 Hz flow oscillations, respectively; and panel e shows data from a human subject during the breath task. (f-h) Summary errors for flow phantom data (panel f, mean error = 0.31 ± 0.21 cm/s); human breath task data at the narrow slice positioning (panel g, mean error = 0.22 ± 0.058 cm/s); and at the wide slice positioning (panel h, mean error = 0.13 ± 0.028 cm/s). Errors were computed by taking the mean absolute difference between measured and predicted velocity, after time-shifting the signals to maximize cross-correlation in order to correct for phase difference arising from separately acquired runs. Predicted velocities were lowpassed below 0.5 Hz to match phase contrast velocity data. (i-j) Relative bandpowers (BP_breath_/BP_total_) defined as bandpower around the paced breathing frequency (0.14-0.19 Hz) divided by the total power, using paced breathing data for fMRI inflow signals (fMRI), phase contrast velocities (PC), and network estimated velocities (NN). Results are shown separately for data at the narrow (panel i) and wide (panel j) slice positioning. Relative bandpowers were significantly different between the phase contrast velocities and estimated velocities at both the narrow (p=0.014; t=-2.7930; paired t-test; Bonferroni adjusted; n=8) and wide (p=0.009; t=-3.4314; paired t-test; Bonferroni adjusted; n=5) slice positions.

To test the accuracy of the network in predicting velocities, we used our measured phantom and human paced breathing fMRI inflow signal data as input, and compared predicted velocities to ground truth velocities measured using phase contrast imaging. We evaluated performance by computing errors as the mean absolute difference between the measured and predicted velocity timeseries. To account for phase differences from separately acquired runs, we time-shifted the signals to maximize cross-correlation, focusing the error calculation on amplitude differences. Representative examples show that the network effectively generalized to infer velocity in real datasets (Fig. 6b-e). Across all runs, the trained network accurately predicted flow velocity using fMRI data and cross-sectional areas for each individual as input (Fig. 6f-h).

We next aimed to quantify whether the predicted flow velocities had high sensitivity to task related changes, as compared to velocities measured using phase contrast imaging. We computed the bandpower around the paced breathing frequency divided by total power in the signal and compared this relative bandpower across flow measured with fMRI, phase contrast imaging, and the network-predicted velocities. Higher values of this relative bandpower metric indicated that the signal contained greater changes in respiratory locked flow relative to noise. We found that the predicted velocities across all paced breathing data had significantly higher relative bandpower than corresponding phase contrast velocities (Fig. 6i,j). The relative power was similar between the fMRI inflow signals and the estimated velocities, suggesting that the high sensitivity provided by fMRI can carry over to the velocity estimations.

### 3.6 Effect of TR and slice thickness on inflow signal amplitude and slice decay rate

The analyses in this study used a model trained using acquisition parameters specific to our dataset. We therefore next aimed to understand the effect of the TR and slice thickness (W) on the inflow effect to estimate the degree of inflow signal expected from varying fMRI protocols. Based on prior work (J. H. Gao et al., 1996; J. H. Gao & Liu, 2012; J.-H. Gao et al., 1988), we expected to observe larger inflow effects for shorter TRs and indeed simulations using constant flow showed that inflow signal amplitude in the first slice decreased with TR (Fig. 7a). Similarly, increasing slice thickness reduced the inflow effect (Fig. 7b), since a larger slice would contain relatively more saturated spins.

**Figure 7.**
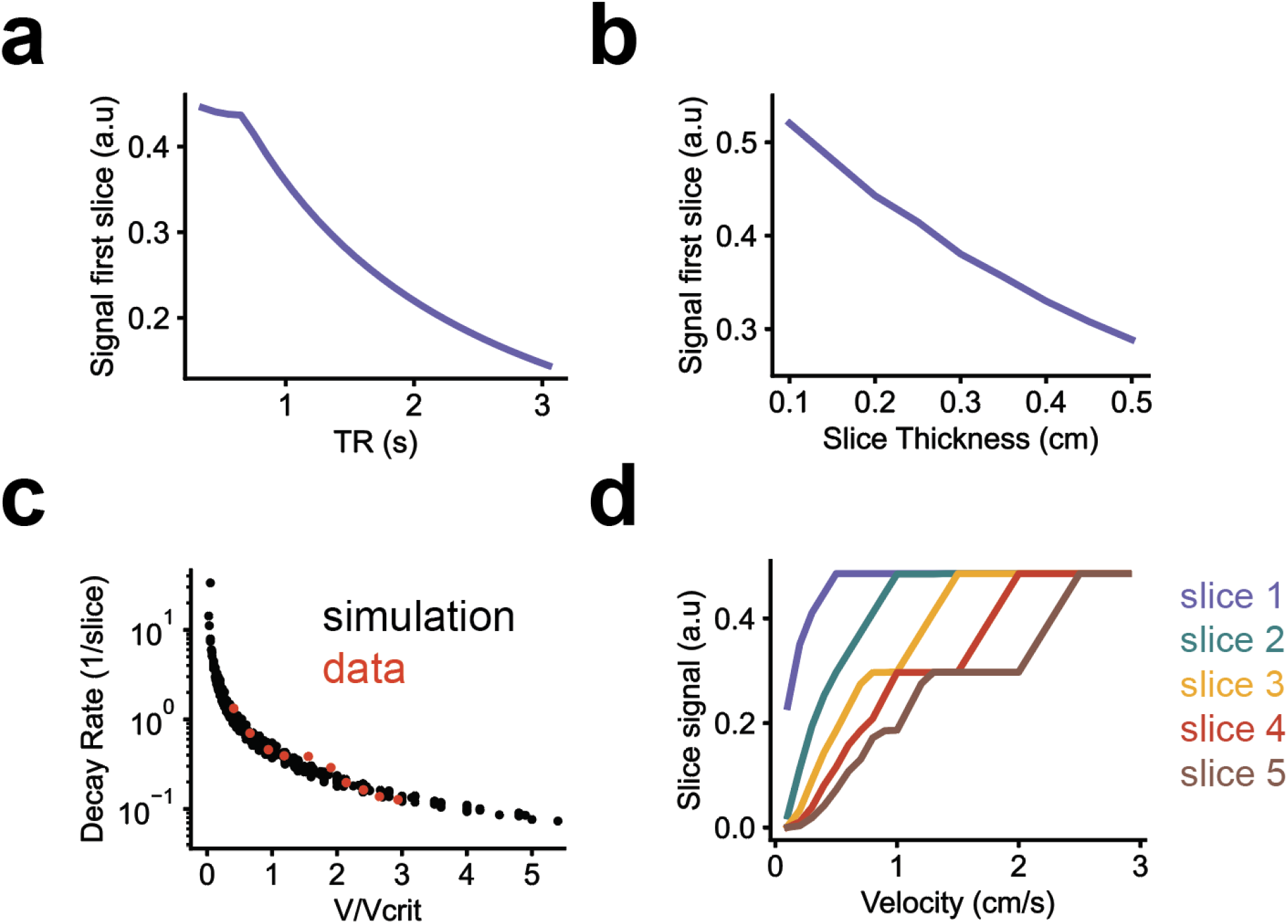
Influence of TR and slice thickness on simulated inflow signal amplitude and slice decay rate. (a) Simulated inflow signal in the first slice as a function of TR. Simulations used constant flow velocity of 0.35 cm/s and a slice thickness of 0.25 cm. Flow was simulated through a straight tube. (b) Simulated inflow signal in the first slice as a function of slice thickness, using same flow input as panel a with a TR of 0.4 s. (c) Signal decay rate across slices as a function of the input velocity divided by the critical velocity (V/V_crit_). Y-axis shown in logscale. Simulations were performed over a range of velocities (0.1-1 cm/s), slice thicknesses (0.1-0.4 cm), and TRs (0.1-0.8 s) and for every combination, decay rate was define as the exponent D of an exponential function (*S*(*x*) = *e*^−*Dx*^) fitted to mean inflow signals (normalized to the first slice) across multiple slices (x = slice index). Measured data during constant flow from the flow phantom is superimposed in red. (d) Simulated inflow signals across five slices simulated using constant flow over a range of velocities (TR=0.4; slice thickness = 0.25) showing saturation of slice signals at each integer multiple of the critical velocity.

In addition to modulating the amplitude of the inflow effect, the TR and slice thickness also influence the signal attenuation across slices, which is where velocity information is encoded and is leveraged by the neural network to estimate velocity. Slower flow results in more decay of signal across the slices due to spins receiving more excitations before moving to the next slice, whereas faster flow results in less decay. How close in time the excitations are sent (TR) as well as the width of the region the signal is sampled from (W) will consequently affect the signal decay across slices. To quantify this relationship, we computed the ‘decay rate’ (defined here as the signal attenuation across slices) of simulated inflow signals under different flow velocities, TRs, and slice thicknesses. For each simulated inflow signal, we fit an exponential function to normalized amplitudes across slices to quantify the decay rate and we analyzed how the decay rate varied with the ratio of input velocity (V) to the critical velocity (V_crit_ = W/TR). Since TR and slice thickness define the relevant velocity scale, this relative velocity measure (V/V_crit_) allowed us to globally assess how the velocity-containing information in the signal varies as a function of a single variable that combines flow velocity, TR, and slice thickness.

Simulation results showed that decay rate decreased exponentially as V/V_crit_ increases and this relationship was consistent with data from constant flow experiments using the flow phantom (Fig. 7c). The sensitivity of velocity estimation from inflow signals is reflected in how much the decay rate changes for a given change in V/V_crit_. We found higher sensitivity when the flow velocity was less than the critical velocity (V/V_crit_ < 1). When V/V_crit_ > 1, signal in the first slice saturates and no longer varies as a function of velocity, reducing degrees of freedom for inferring velocity. However, this does not mean that information about flow speed is completely lost, as signals in deeper slices can still vary relative to each other. As the velocity exceeds integer multiples of the critical velocity, more slices become saturated (e.g. second slice saturates at 2*V_crit_), further reducing velocity-dependent signal variations (Fig. 7d).

## 4. Discussion

This work developed a framework linking fluid flow velocities to fMRI signals, enabling quantification of CSF flow dynamics from fMRI data. We first developed a forward model to capture the relationship between time-dependent flow velocity and the observed fMRI inflow signals. The model accurately captured the dynamics of fMRI inflow signals during constant and oscillatory flow, while accounting for individual anatomy. We then developed an inverse model using a neural network, enabling direct quantification of fluid flow velocities from measured fMRI inflow signals. This work opens an avenue for future neuroimaging studies of dynamic flow using fMRI, enabling quantitative and temporally specific investigations of CSF flow dynamics in fMRI data.

Our modeling results identify key characteristics of the fMRI inflow signal that should be taken into account when analyzing CSF flow in fMRI datasets. Both our data and simulations demonstrate that the frequency spectrum of the inflow signal is affected by a frequency-dependent bias, where the fMRI signal power is attenuated for faster fluid flow oscillations. The level of attenuation for speed fluctuations at a given frequency depends on its distance from the Nyquist frequency. If a brief fluctuation in flow speed (e.g., due to a cardiac pulse) occurs that causes flowing fluid to momentarily slow down and speed back up over a time period shorter than the TR, then the net displacement of the fluid from that flow fluctuation can be small by the time the next RF pulse is received. This results in less fMRI inflow signal power from that oscillatory frequency. In contrast, prolonged fluctuations in flow speed are sampled more as more RF excitations occur over the full oscillation period and therefore its effect on the signal is stronger. Therefore, the measured signal amplitude will decrease as the frequency of oscillations approaches the imaging Nyquist rate. This frequency attenuation effect (Fig. 5) suggests that the interpretation of frequency spectra from fMRI inflow signals is not straightforward. For example, this effect could potentially lead to underestimation of flow contributions from the cardiac cycle, a known modulator of CSF flow dynamics (Mestre et al., 2018), since those frequencies would be strongly attenuated. This confound suggests that modeling will be needed to compare spectral information in fMRI inflow signals across studies, particularly because different studies have different temporal resolution.

We also show that slice placement and individual anatomy is important to consider when imaging flow using fMRI (Fig. 4), since the flow velocity varies with cross-sectional area which in turn affects the magnitude of fMRI inflow signal. The implications of this volume-dependent bias is two-fold: 1) comparing signals measured at locations with different cross-sectional area can introduce flow differences purely from anatomy (Fig. 4c-g), and 2) it is important to account for the spatial dependence of the input velocity when simulating inflow signals (Fig. 4h-j). We recommend that researchers follow a consistent convention when positioning imaging slices in their experiments, aiming to image the narrowest region that avoids partial voluming, in order to obtain higher signal (due to imaging at a location with higher velocity). We also recommend positioning the edge slice such that multiple slices intersect the flow compartment, since the model uses information about relative signal intensities across slices to quantify flow velocity. Positioning the edge of the acquisition to align with the base of the fourth ventricle can achieve both goals. Additionally, our neural network takes cross-sectional areas as input so anatomical imaging should be performed in order to extract that information from each individual. Lastly, it is recommended to prioritize using shorter TRs as that will generally increase inflow-weighting, increase sensitivity of the velocity estimation (Fig. 7f), and reduce the frequency attenuation effect (although we note that inflow has successfully been detected with longer TRs as well (Gonzalez-Castillo et al., 2022; Han et al., 2021; Williams et al., 2023)). Despite these characteristics of the fMRI inflow signal, our physics-based inverse approach sidesteps both frequency-dependent and volume-dependent confounds, allowing researchers to obtain physically interpretable information from fMRI inflow signals and analyze flow velocity directly.

Phase contrast imaging is a gold standard technique for quantitative flow imaging in MRI, and has been used extensively to image blood flow and more recently large-scale CSF flow^8,25^. Phase contrast has high clinical utility and offers distinct information relative to fMRI, which is widely available to neuroscientists and has different sensitivity properties. Due to these distinct properties, fMRI could become a complementary technique to phase contrast, which has classically had relatively poor sensitivity to slow flow, although impressive recent advancements in gradient technology could now enable expanded use cases for phase contrast (Williamson et al., 2020). A key advantage of using fMRI for flow imaging is the simultaneous measurement of BOLD signals alongside the inflow signals which can provide valuable information about how concomitant changes in hemodynamics drive CSF flow. Ultimately, ultra-high field fMRI may be an important future direction for CSF flow imaging because of its high sensitivity, simultaneous BOLD signals, and improved ability to image small spaces due to its high spatiotemporal resolution (De Martino et al., 2018; Dumoulin et al., 2018; Feinberg et al., 2023). Future studies may choose to use ultra-high field fMRI to image fluid flows with simultaneous hemodynamic measurements, and would benefit from adopting our modeling framework to enable non-invasive flow velocity quantification at fast timescales and small spatial scales.

An important limitation in the forward model is that we assume the RF pulse slice selection profile for each slice is a step function covering the full slice. In reality, the flip angle varies across the slice, and this property can affect flow measurements (Zong & Lin, 2019). Future work could modify the flip angle parameter to be a function of position within the slices to enable different options for the slice selection profile. Additionally, the scan parameters of our specific fMRI protocol were fixed during training of the neural network. Future studies applying this modeling framework can tailor the scan parameters to their specific imaging protocols during training to obtain a specialized inverse model for their dataset. Furthermore, our simulated dataset used for training the neural network assumed a specific structure for CSF flow oscillations based on dynamics observed in our human data (see Methods), but future studies could expand the nature of dynamics included in the network. Our code is shared openly to enable further customization based on the parameter settings, physiological dynamics, and image noise properties of local datasets across labs. We could not examine cardiac oscillations using our neural network approach due to the limited temporal resolution of our phase contrast protocol. We focused here on validating performance of slower oscillations and prioritized maximizing SNR in the 2.5mm slice over temporal resolution for the phase contrast protocol. Phase contrast imaging with higher temporal resolution will be necessary in future work to validate the estimations of faster oscillations. Furthermore, our implementation of the spatial dependence of velocity assumed that velocity scales inversely with area in the fourth ventricle, which neglects complex flow patterns that could emerge. The fMRI flow measurement approach is limited in that it measures flow orthogonal to the imaging slice, and can only quantify flow in that direction; understanding more complex spatial properties of flow patterns requires different techniques such as 4D phase contrast imaging in the 4^th^ ventricle (Rivera-Rivera et al., 2024).

This framework could next be broadly applied to investigate CSF dynamics in the human brain. Due to the wide availability of large public fMRI datasets, including the HCP dataset which is known to contain CSF signals (Gonzalez-Castillo et al., 2022), as well as the wide availability and use of fMRI, this framework would allow CSF analyses in diverse patient samples. Furthermore, due to its incorporation of individual anatomy, this framework is well suited to examine individual differences, such as studying individuals across the human lifespan, when ventricles expand substantially. It can also be readily applied to other CSF-containing regions simply by changing the acquisition positioning of the fMRI volume. Finally, this model can also benefit future experimental designs by providing the ability to simulate how different fMRI protocol parameters (such as the TR, slice excitation timings, and slice thickness) affect the amplitude of distinct types of flow signals, which will be beneficial for designing future fMRI studies seeking to measure flow with high sensitivity.

## 5 Conclusion

The dynamics of CSF flow are an essential component of the biological systems that maintain healthy brain function. fMRI has been used to measure CSF flow in the human brain, providing a useful method for monitoring the flow dynamics with high sensitivity and temporal resolution, as well as unique information about hemodynamics. We present a framework for inferring quantitative information about fluid flow velocities from non-quantitative fMRI data. This work advances the theoretical understanding of flow-enhanced signals caused by dynamic flow, and provides a practical approach that neuroimaging researchers can use to study fluid flow using fMRI.

## Ethics

All human participants gave informed consent before participating in this study. This study was approved by the Boston University Institutional Review Board.

## Data and Code availability

Data will be shared publicly upon manuscript acceptance. Code is publicly available at https://github.com/baarbod.

## Author Contributions

B.A., L.D.L, and D.E.P.G developed the model. B.A., and D.E.P.G collected data. B.A., L.D.L., and D.E.P.G wrote the manuscript.

## Acknowledgements

We are grateful to Amelia Strom, Zijing Dong, Jonathan Polimeni, and Makaila Banks for useful conversations and assistance with data acquisition. We thank Bastien Guerin for providing a prototype flow phantom.

## Materials & Correspondence

Correspondence to Laura D. Lewis.

## Appendix

Let *C* = cos *θ* and 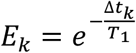, where θ is the RF excitation flip angle, T_1_ is the longitudinal relaxation time constant, and Δ*t*_*k*_ is the amount of time between the excitation pulse k and k+1. Recall that the longitudinal magnetization (M_z_) is described from the Bloch equations(Bloch, 1946) by

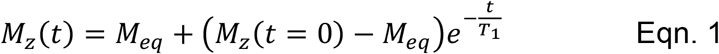

which describes the exponential relaxation of M_z_ back to equilibrium state M_eq_ from an initial magnetization *M*_*z*_(*t* = 0) after excitation by an RF pulse. The longitudinal magnetization at equilibrium just before (indicated by the − superscript) the first excitation is

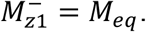

An excitation scales M_z_ by C so that the longitudinal magnetization just after the excitation (indicated by the + superscript) becomes

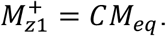

Substituting the above expression for M_z0_ in Eqn. 1 and setting t equal to Δ*t*_1_, we obtain

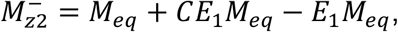

which is the longitudinal magnetization after a time period Δt_1_ of relaxation between the first and second excitations. Immediately after the second excitation, M_z_ is scaled again by C and we thus obtain M_z_ just after the second excitation by multiplying the above expression by C

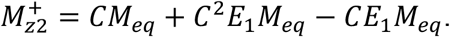

This process is repeated for all subsequent excitations received. We show below this process up to the moment just before the fourth excitation.

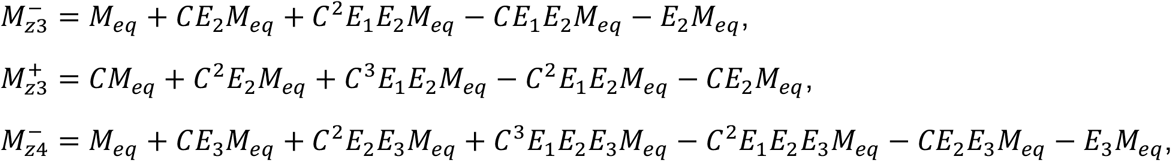

We can derive a general expression by grouping the positive and negative terms each into separate summation series, and expressing the exponential terms as ratios of product series. The general expression as a function of the number of excitations (n) and all the time-periods (Δt) between each excitation leading up to the n^th^ excitation is

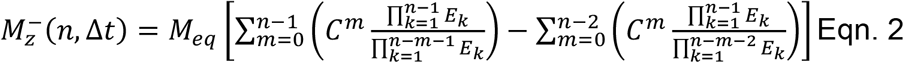

We can show an equivalence between this new expression and a prior one(Carr, 2012) that assumed all inter-pulse times were TR by setting all Δt’s in Eqn. 2 equal to TR.

Making this substitution and renaming E_k_ to E (since all E_k_ will then be identical) allows us to simplify the two inner terms of each series in Eqn. 2 as follows

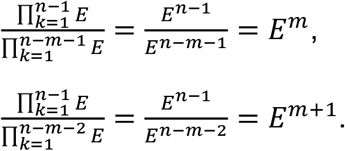

Substituting these simplified expressions back into Eqn. 2 and simplifying yields

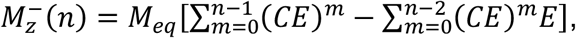

where we can apply a finite summation identity to obtain

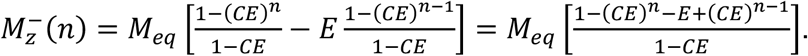

At steady state, the number of excitations n approaches infinity resulting in (*CE*)^*n*^→ 0, because the inner term is less than 1. Thus, we obtain the following equation at steady state which is equivalent to the prior expression (equation 2.3 in reference(Carr, 2012))

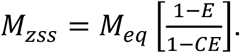

This equivalence shows that we have generalized the prior relationship, enabling us to model the effect of variable time periods between excitations. We can further simplify Eqn. 2 by modifying the upper bound of one of the series as shown below such that the two remaining series have the same upper bound and could be combined.

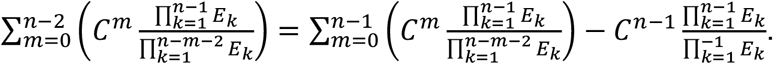

Plugging the above expression back into Eqn. 2 and combining the two series yields

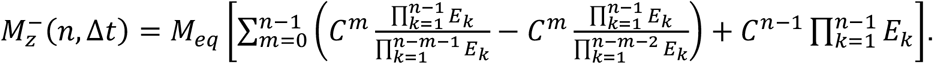

After factoring we obtain

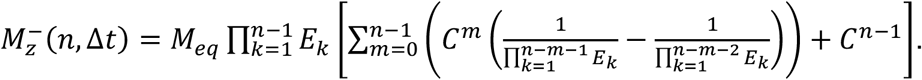

The final expression used in the programming implementation after combining the two fractions in above expression and converting the longitudinal magnetization to the transverse magnetization (M_T_) is the following

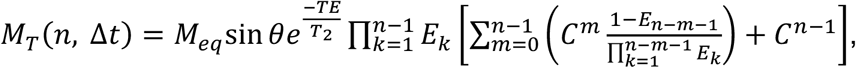

where TE is the echo time, and T_2_ is the transverse relaxation time constant.

