## Supplemental Materials for "Modeling dynamic inflow effects in fMRI to quantify cerebrospinal fluid flow"

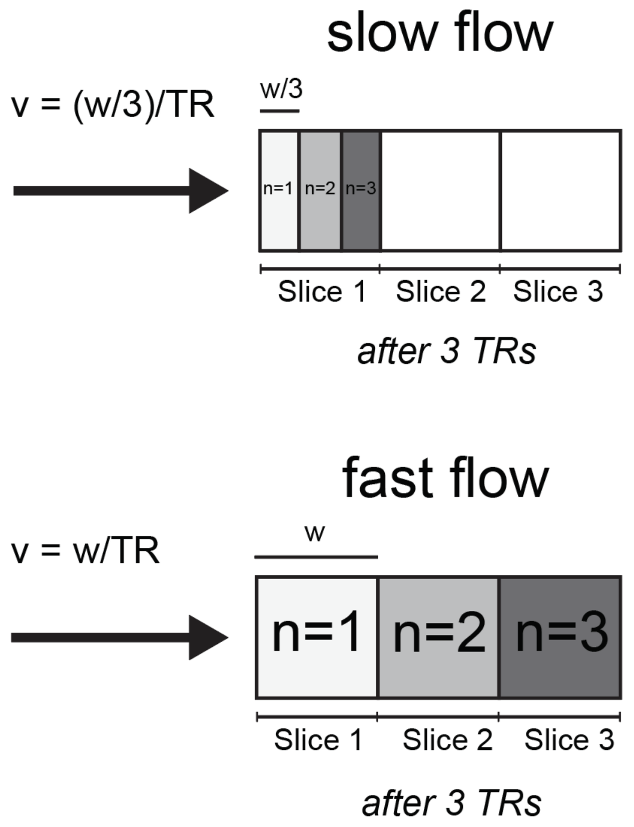


**Figure S1. Example scenario with plug flow showing how flow speed affects how fluid is excited across imaging slices.** Illustration using constant plug flow of how fluid flow speed affects the number of excitations received by fluid in three slices after three repetition cycles. Slow flow results in more excitations accumulated in the first slice, in contrast to fast flow which results in a fully saturated first slice.

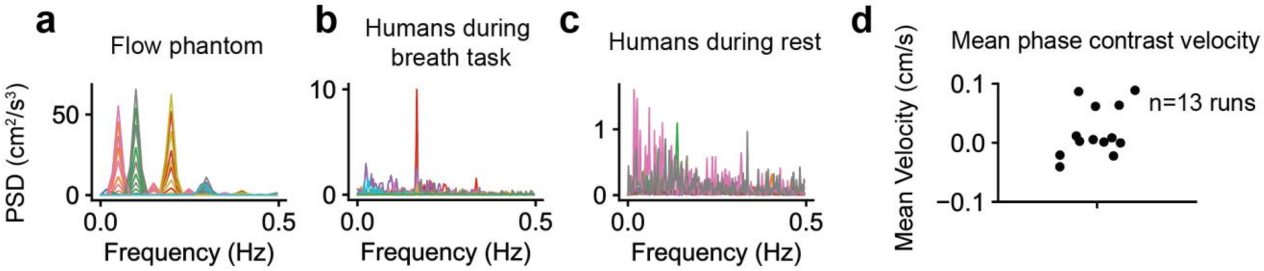


**Figure S2. Measured velocity spectra from phase contrast imaging.** In humans, power spectral density (PSD) was calculated from the phase contrast velocity signal of CSF flow in the fourth ventricle, which typically shows oscillatory dynamics locked to breathing. (a-c) Overlayed frequency spectra from experiments with the flow phantom system (a; 30 runs), humans during a 0.167 Hz paced breathing task (b; 13 runs), and humans during resting state (c; 10 runs). Distinct colors represent data from distinct runs. (d) Mean phase contrast imaging velocities for all runs which was used to inform velocity sampling bounds used for the inverse model. Each dot represents the mean value of the velocity timeseries.

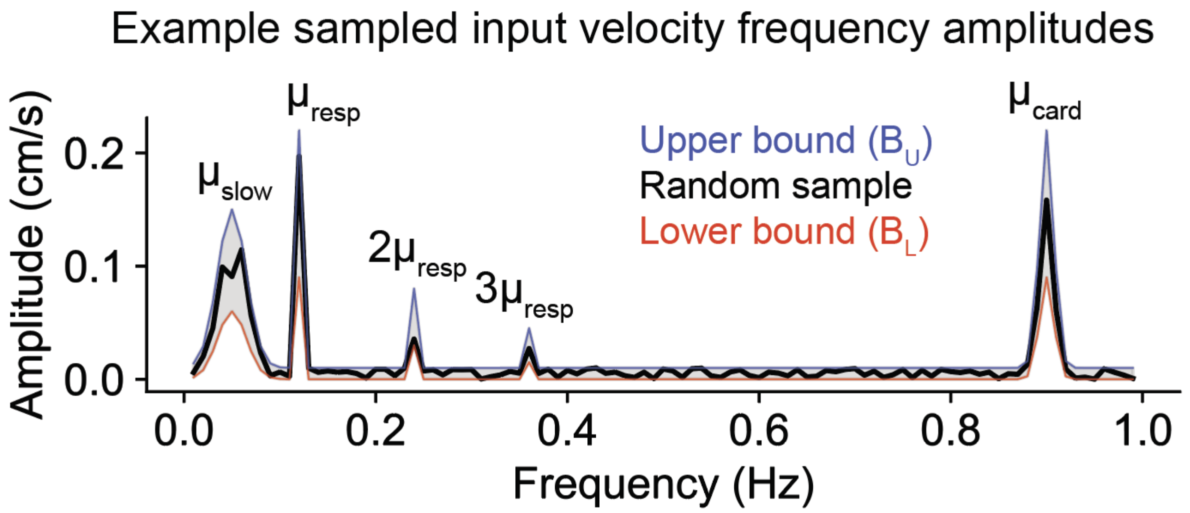


**Figure S3. Example sample distribution for simulating input velocity to generate training data, with frequency amplitudes randomly sampled between lower and upper bound curves.** The lower (red) and upper (blue) curves represent frequency-dependent sampling intervals used for uniform random sampling. Taking a random value for each frequency, within the bounds, yields frequency amplitudes for a sample input velocity used in the training dataset (black). This example used $\alpha_{slow}$ = 0.1 cm/s, $\mu_{slow}$ = 0.05 Hz, $\alpha_{resp}$ = 0.15 cm/s, $\mu_{resp}$ = 0.12 Hz, $\alpha_{card}$ = 0.15 cm/s, and $\mu_{card}$ = 0.9 Hz. This example illustrates the sampling distribution for a particular sample; during training these parameters vary.

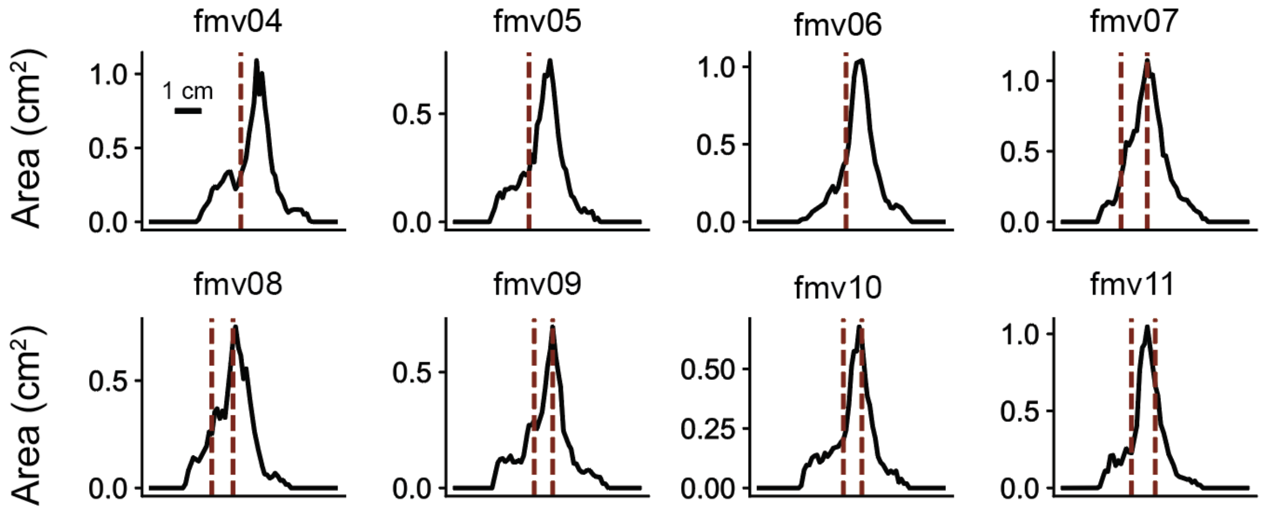


**Figure S4. Slice positioning for all human subjects.** Cross-sectional area curves extracted from individual anatomical images are shown with overlayed red vertical dashed lines indicating the locations where the edge of the bottom fMRI slice was positioned. The first three subjects (fmv04, fmv05, and fmv06) only had a single slice positioning within the experiment, while the subsequent five subjects had data from two different slice positioning: one at a narrower part of the fourth ventricle and another at a wider part. Scale bar on the top left panel is shown for a 1 cm distance.

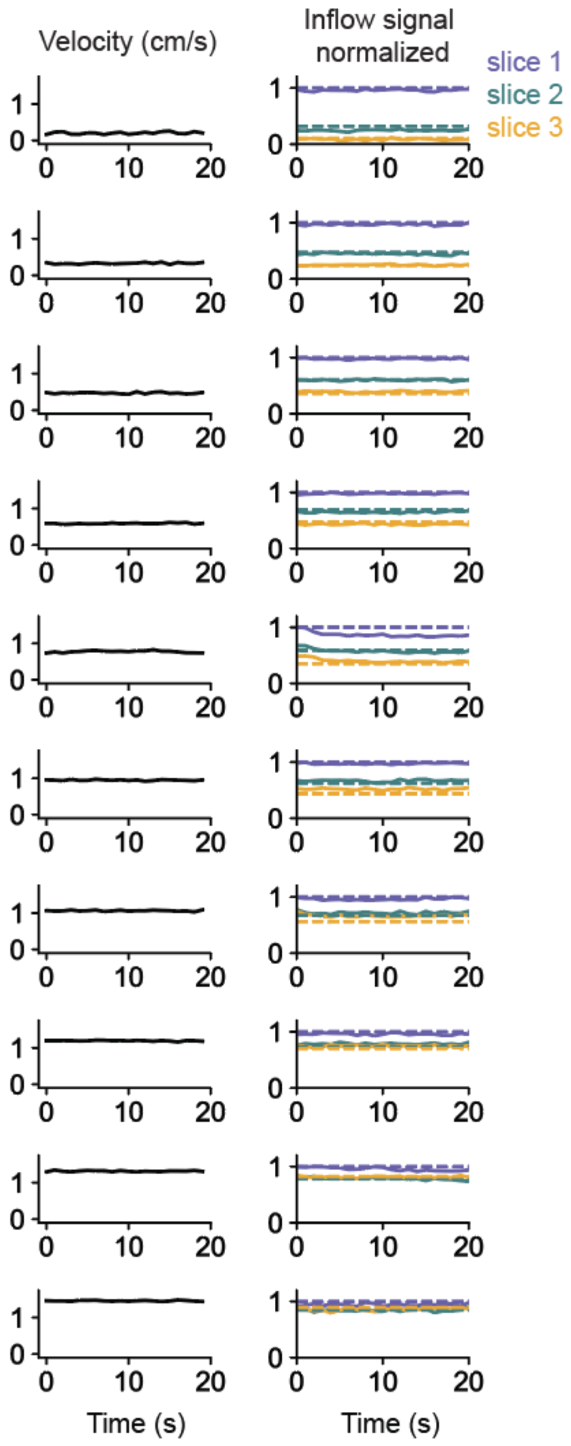


**Figure S5. Model performance in a flow phantom using constant velocities.** Left column: Phase contrast measured velocities tested in order of increasing speed; Right column: corresponding measured fMRI inflow signals (solid lines) in the first three slices overlayed with simulated fMRI inflow signals (dashed lines) using the mean velocity of each phase contrast trace as a constant flow input into the model. All inflow signals are normalized to the mean value of the top 5% of signals in the first slice.

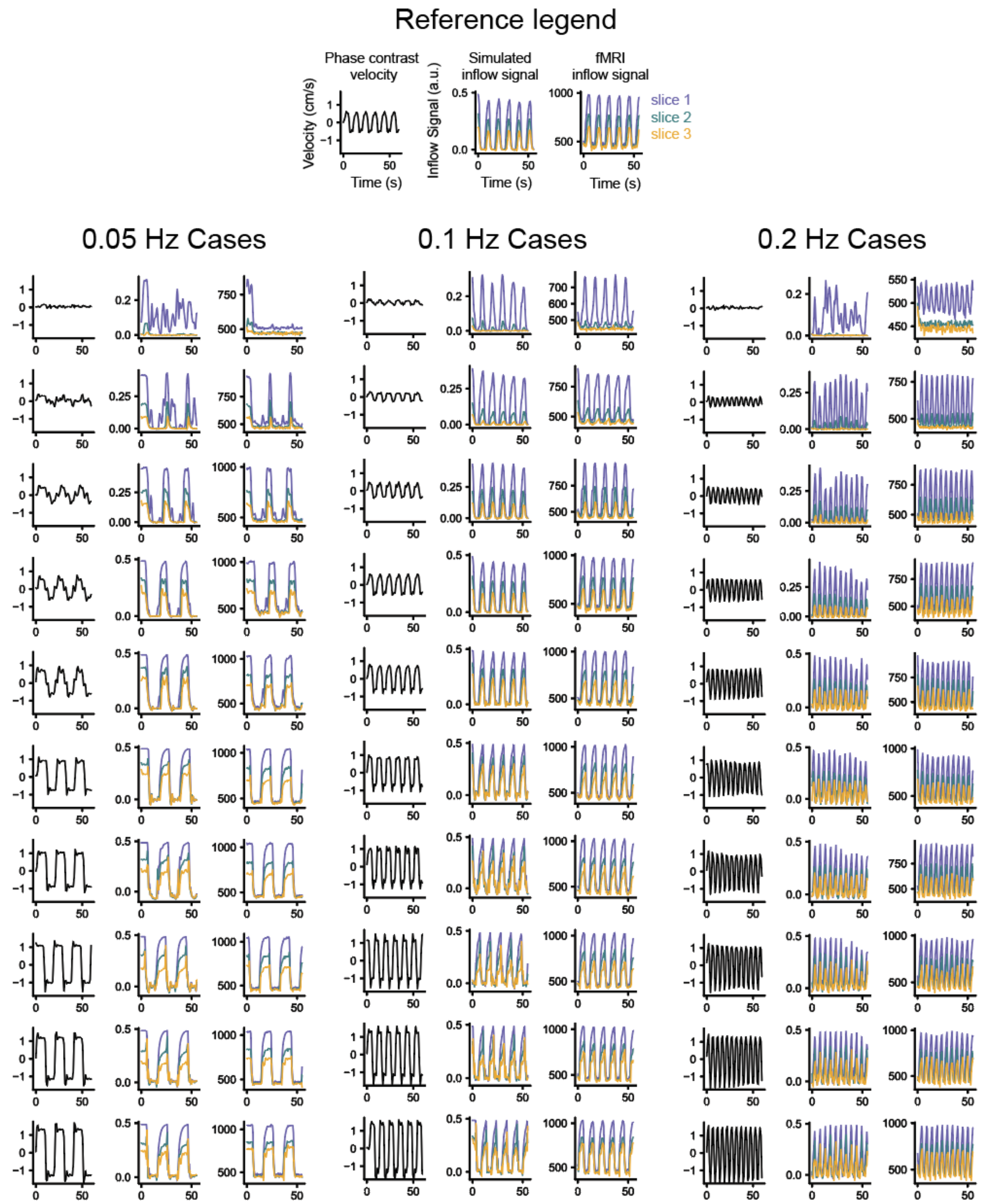


**Figure S6. Model performance in a flow phantom using time-varying velocities.** Left group), left column: phase contrast measured velocities oscillating at 0.05 Hz; middle column: corresponding measured fMRI inflow signals in the first three slices; right column simulated fMRI inflow signals using phase contrast velocities as input into the model. The middle and right group show data for 0.1 Hz and 0.2 Hz oscillations. All inflow signals are normalized to the mean value of the top 5% of signals in the first slice.

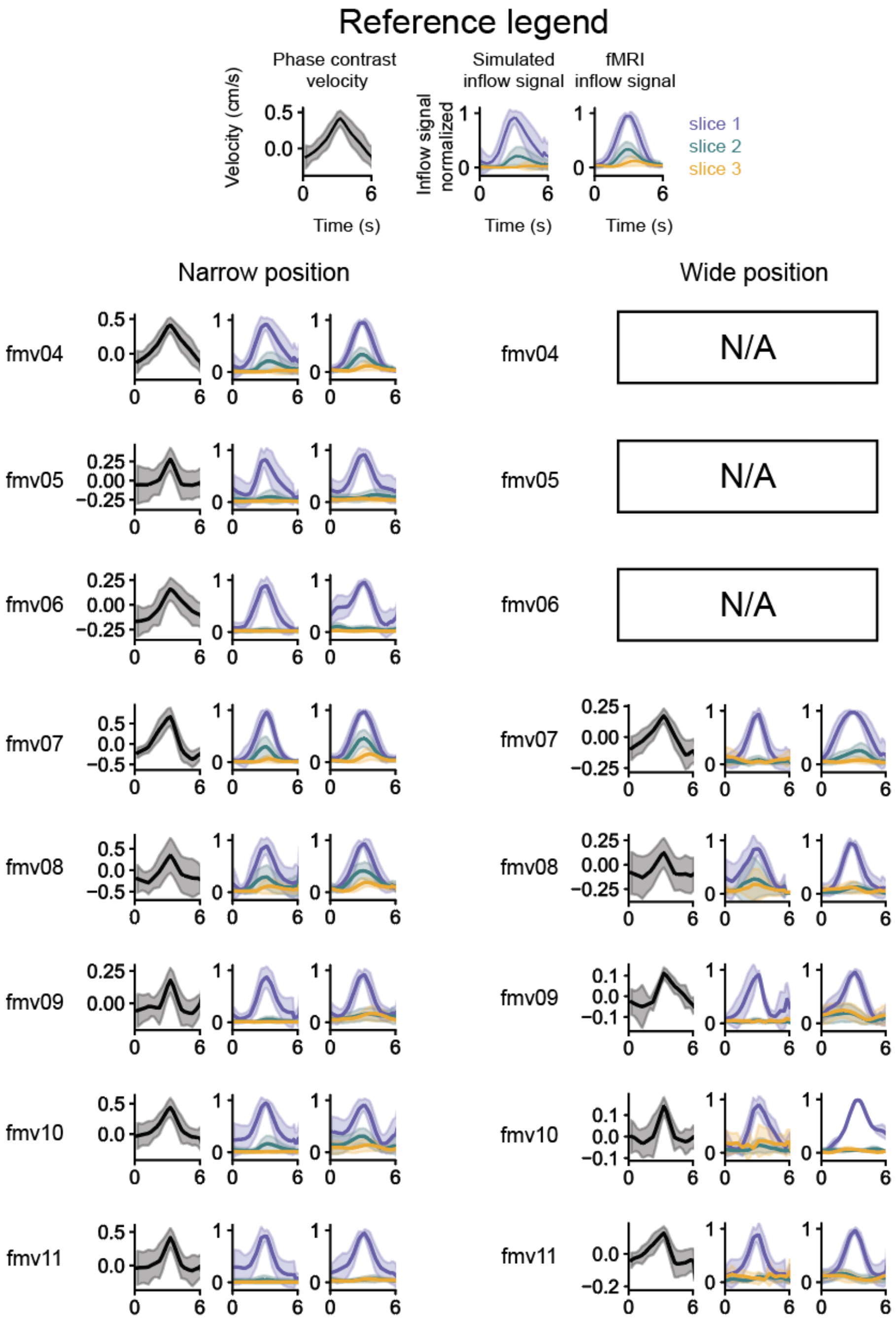


**Figure S7. Period-averaged human data and simulations during paced breathing.** Data is shown from the narrow (left group) and wide (right group) slice positionings. The first three subjects (fmv04, fmv05, and fmv06) only had a single slice positioning within the experiment, while the subsequent five subjects had data from two different slice positioning. Each group of three panels shows period-averaged data for each subject: phase contrast velocity, simulated inflow signal using the velocity as model input, and measured fMRI inflow signals (see reference legend). Data shown is during 0.167 Hz paced breathing. All inflow signals are normalized to the mean value of the top 5% of signals in the first slice, after subtracting.
